# Selective scoring of drug effects in multicellular co-culture systems

**DOI:** 10.64898/2026.05.20.726737

**Authors:** Diogo Dias, Aleksandr Ianevski, Jonas Bouhlal, Michele Ciboddo, Petra Nygrén, Jay Klievink, Hanna Lähteenmäki, Olli Dufva, Satu Mustjoki, Tero Aittokallio

**Affiliations:** Institute for Molecular Medicine Finland (FIMM), HiLIFE, University of Helsinki, Helsinki, Finland; iCAN Digital Precision Cancer Medicine Flagship, University of Helsinki and Helsinki University Hospital, Finland; Hematology Research Unit Helsinki, University of Helsinki and Helsinki University Hospital Comprehensive Cancer Center, Helsinki, Finland; Translational Immunology Research Program and Department of Clinical Chemistry and Hematology, University of Helsinki, Helsinki, Finland; Institute for Cancer Research, Department of Cancer Genetics, Oslo University Hospital, Oslo, Norway; Oslo Centre for Biostatistics and Epidemiology (OCBE), Faculty of Medicine, University of Oslo, Oslo, Norway; Cambridge Stem Cell Institute, Department of Medicine, University of Cambridge, Cambridge, United Kingdom; Department of Cellular Genetics, Wellcome Sanger Institute, Hinxton, United Kingdom

**Author notes:** Shared senior authors and correspondence: Tero Aittokallio, PhD, Institute for Molecular Medicine Finland (FIMM), P.O. Box 20 (Tukholmankatu 8), FI-00014 University of Helsinki, Helsinki, Finland. Satu Mustjoki, MD, PhD, Hematology Research Unit Helsinki, University of Helsinki and Helsinki University Hospital Comprehensive Cancer Center, Haartmaninkatu 8, FI-00290 Helsinki, Finland. Equal contribution as second authors (Aleksandr Ianevski, Jonas Bouhlal).

## Abstract

Multicellular co-culture screening reveals compound effects that depend on cell-cell interactions. Standard dose-response metrics fail to resolve effects that arise either from target-effector cell interactions or from non-specific toxic effects. Here, we developed Co-culture Efficacy Score (CES), a robust computational framework that enables systematic identification of compounds that selectively modulate cellular interactions in multicellular assays. CES framework supports both therapeutic scoring that penalizes direct effector cell toxicity, as well as a mechanistic discovery that estimates immunomodulatory effects by adjusting for effector cell responses. When screening 527 compounds across 10 hematological cancer models co-cultured with natural killer (NK) cells, CES distinguished co-culture-specific immunomodulatory effects from NK cell toxicity and cancer cell inhibitory responses, recovering systematic enhancer and inhibitory patterns. We further assessed CES robustness using higher-resolution validation screens and demonstrated its applicability to identify selective compounds in anti-CD19 CAR T-cell and antiviral host-pathogen screens. To facilitate its broad use, we implemented CES as an interactive web-application for quantitative analysis of compound responses in co-culture assays, providing a widely applicable scoring framework for cancer immunotherapy, antiviral screening and drug discovery.

## Introduction

Cell-based compound screening provides a powerful functional approach to systematically interrogate pathway activity and cell signaling, as well as to identify therapeutic vulnerabilities in cancer cells. The impact of these high-throughput assays relies on analytical methods that quantify compound activity across diverse experimental conditions. Such quantitative pharmacological profiling approaches have critically supported compound mode-of-action discovery and functional precision medicine strategies, where direct testing of drug activity in patient-derived cells informs treatment optimization and supports clinical decision making^1-10^.

Most large-scale compound sensitivity screens to date have tested compounds in cancer cell models under monoculture conditions. These simplified assays fail to capture the full spectrum of compound effects that arise from cellular perturbations within complex biological systems^11^. Since compound activity is strongly influenced by interactions between target tumor cells and immune effector cells, or other components of the tumor microenvironment, there is a strong motivation to develop co-culture assays that capture these complex cell-cell interactions and context-dependent activities both in hematological malignancies and solid tumors^12-19^. Conceptually similar assay setups arise also in non-cancer screening contexts, including antiviral screens, where host cells are cultured either alone with a compound or together with a viral pathogen in a co-culture format, and a selective compound activity is quantified in terms of whether the treatment rescues host-cell viability from virus-induced cytopathic effects^20-23^.

Cancer immunotherapy has further triggered the need for more complex experimental systems capable of capturing interactions between immune effector cells and tumor target cells^24^.

Natural killer cells^25,26^ and cytotoxic T lymphocytes^27^ are key mediators of anti-tumor immunity, directly eliminating malignant cells through diverse cytotoxic and immunoregulatory mechanisms, yet these responses can be further enhanced or suppressed by targeted pharmacological perturbation. Co-culture screening assays provide an attractive strategy for identifying compounds that modulate effector-cell activity and tumor susceptibility in combinatorial therapy settings. Integrated drug profiling and functional genomic screens have enabled systematic investigation of mechanisms regulating CAR T-cell cytotoxicity and T-cell effector function^28-30^, while pharmacological perturbation screens have shown that drug treatment reprograms NK cell-tumor interactions and therefore enhances NK cell-mediated cytotoxicity^31,32^.

However, existing dose-response metrics used in large-scale screening have been developed for single responding cell populations under monoculture conditions. Dose-response modeling in cell-based assays typically relies on parameters, such as the half-maximal inhibitory concentration (IC_50_), and minimum and maximum response levels^33,34^. These parameters form the basis for summary response metrics, including the area under the curve (AUC) and drug sensitivity scores (DSS)^35^, which are widely used to quantify compound responses in diverse applications^36-40^. Such scoring frameworks have further been applied to quantify selective drug responses relative to healthy cells to guide personalized treatment strategies^41^. A central challenge posed by the co-culture systems is that these metrics assume a single target cell population (e.g., tumor cells), and therefore fail to resolve effects arising from the interactions with effector cells (e.g., immune cells) and chemical perturbations.

We developed the Co-culture Efficacy Score (CES), a model-based analytical framework for robust analysis and prioritization of compound responses in multicellular screens using paired co-culture and monoculture measurements. CES models condition-specific dose-response relationships through integrated response profiles that capture modulation by effector-mediated cytotoxicity relative to the monoculture conditions. Due to non-monotonic response trajectories, CES uses Gaussian mixture modeling to estimate response parameters that distinguish selective effector modulation from non-specific toxicity, hence prioritizing compounds that enhance or inhibit co-culture responses. CES supports two complementary scoring applications: (i) a therapeutic selectivity through penalizing effector-cell toxicity via the Bliss independence model, and (ii) a mechanistic discovery by adjusting the interaction signal by the surviving effector cell fraction.

Here, we evaluated CES in large-scale NK-cell co-culture screens, higher-resolution validation experiments, CAR T-cell co-culture assays, and antiviral host-pathogen applications. Across these diverse applications, CES identified compounds that selectively enhanced or inhibited cell-cell interaction phenotypes, improved interpretability relative to conventional dose-response metrics, including IC_50_, AUC, DSS, therapeutic index and selectivity index, and provided a reproducible framework for prioritizing compounds in multicellular screening assays. To support its broad adoption, we implemented the CES framework as a user-friendly, interactive and fast web-application for streamlined analysis of co-culture screening datasets.

## Results

### A computational framework for integrated dose-response modeling in co-culture compound screening

Modeling compound responses in co-culture systems is challenging because the measured phenotype reflects contributions from more than one cell population, which generates response patterns that differ substantially from standard monoculture dose-response curves^11^. To address this challenge, we developed CES, a robust computational framework that quantifies context-dependent drug activity by comparing co-culture responses with matched reference conditions. The CES framework enables compound prioritization across multiple cell models and experimental conditions (**Fig. 1a**). Each cellular model is typically evaluated under three conditions: (i) drug treatment in the target cell monoculture, (ii) the corresponding co-culture with effector cells, and (iii) assessment of drug-induced toxicity in the effector cell population. In studies where the full three-condition profiling is not available, CES supports two-condition applications, utilizing either target monoculture or effector toxicity measurements alongside the co-culture responses.

**Figure 1.**
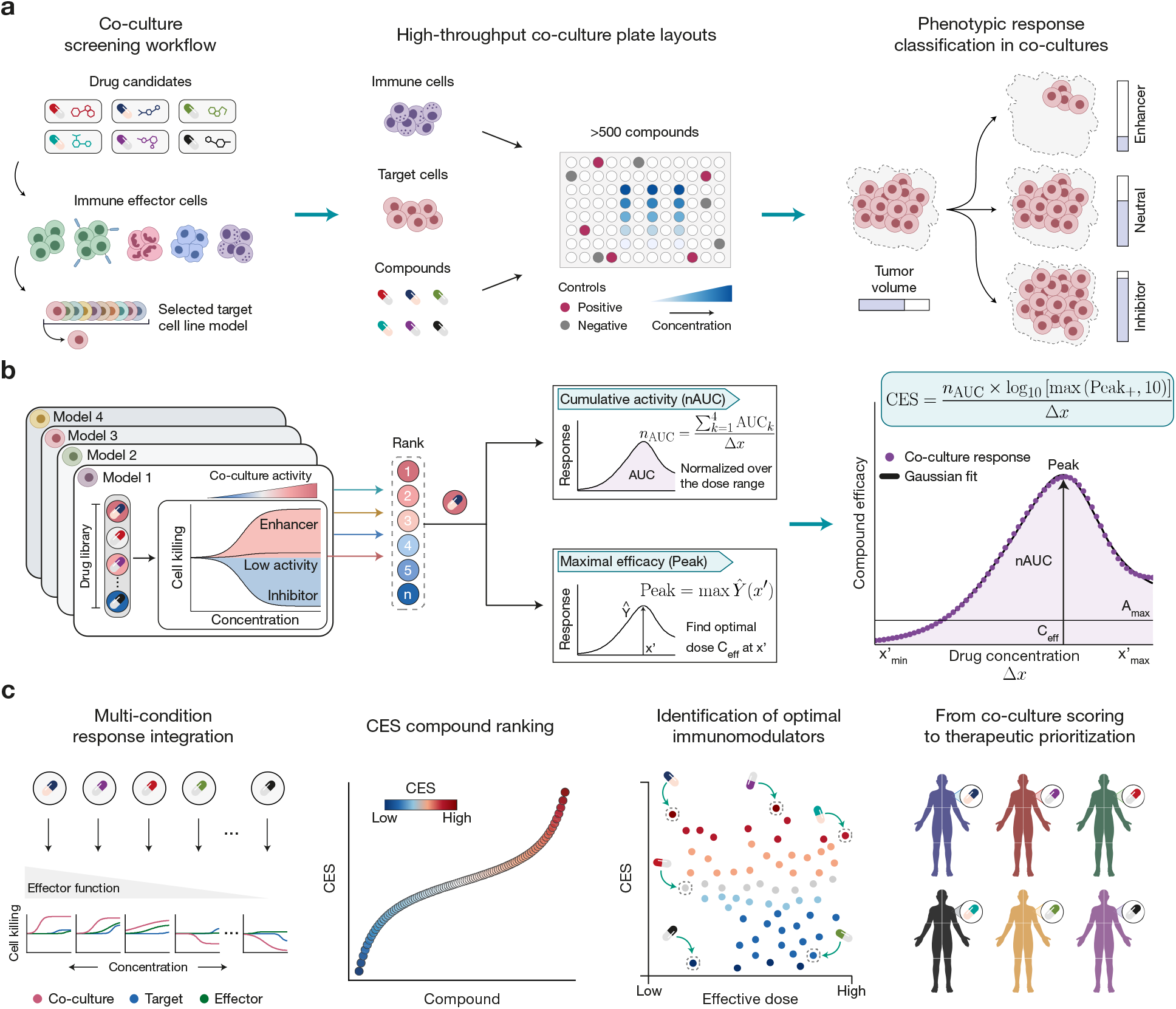
Overview of the CES framework for co-culture drug response quantification. (**a**) A compound library is screened across multiple cancer cell models paired with immune effector cells or other interacting cell types. For each model, three parallel experimental conditions are evaluated across a defined concentration range: (i) target cell monoculture, (ii) target-effector co-culture, and (iii) effector cell monoculture. Compounds are categorized as enhancers, inhibitors, or neutral agents based on their modulation of effector-mediated cytotoxicity. (**b**) Condition-specific dose-response curves are combined into an integrated co-culture response profile, which is fitted using Gaussian mixture modeling. Profiles with insufficient activity (A_max_ < 10% by default) are assigned CES = 0. The fitted model is used to extract cumulative activity (nAUC), maximal effect (Peak), and effective dose (C_eff_) parameters. These quantities are integrated to compute the Co-culture Efficacy Score (CES), which enables standardized comparison of compound activity across experimental models. (**c**) Integrated co-culture response profiles from all conditions are used for CES-based compound ranking, visually distinguishing top enhancers (red) from strong inhibitors (blue). Ranked compounds are then mapped against their effective doses to identify model-estimated effective concentrations for modulating effector-mediated responses. Quantitative scoring and compound prioritization support selection of context-dependent modulators across diverse therapeutic applications.

To capture non-monotonic or bell-shaped integrated co-culture response profiles derived from multiple screening conditions, CES makes use of Gaussian mixture modeling, rather than relying on standard sigmoidal fits (**Fig. 1b**). From these fitted dose-response models, CES estimates core quantitative parameters, including cumulative activity, quantified as normalized area under the response curve (nAUC), maximal effect (Peak), and effective dose metrics. These features are integrated into a single metric that ranks compounds based on the magnitude and consistency of their co-culture-specific effects (**Fig. 1c**). Intuitively, positive CES values indicate an enhanced effector-mediated killing beyond direct toxicity to the target cells, whereas negative values indicate inhibition of effector-mediated killing, with higher magnitudes in either direction reflecting more pronounced effects. By combining broad activity across the tested dose range with maximal response magnitude, CES prioritizes compounds with robust context-dependent efficacy, rather than relying on a single potency estimate.

### CES prioritizes interaction-selective compounds in NK-cell co-culture screens

We first evaluated CES using our previously generated NK-cell co-culture compound screening dataset^31^, comprising ten hematological cancer cell line models profiled against 527 compounds. A total of 15,800 dose-response profiles were analyzed across target-NK cell co-culture, target monoculture, and effector monoculture conditions (**Supplementary Data 1**). The resulting dataset captured substantial heterogeneity in compound sensitivity, reflecting diverse genetic backgrounds and context-dependent immune modulation (**Fig. 2a**). Plate-level quality control based on positive and negative control wells demonstrated robust assay performance across 184 screening plates, with a median Z′ of 0.81 (IQR 0.66–0.88); 92.4% of plates exceeded the standard Z′ threshold of 0.5. Robust Z′ values showed similar performance (median 0.80; IQR 0.64–0.89) (**Supplementary Fig. 2a**; **Supplementary Data 2**).

**Figure 2.**
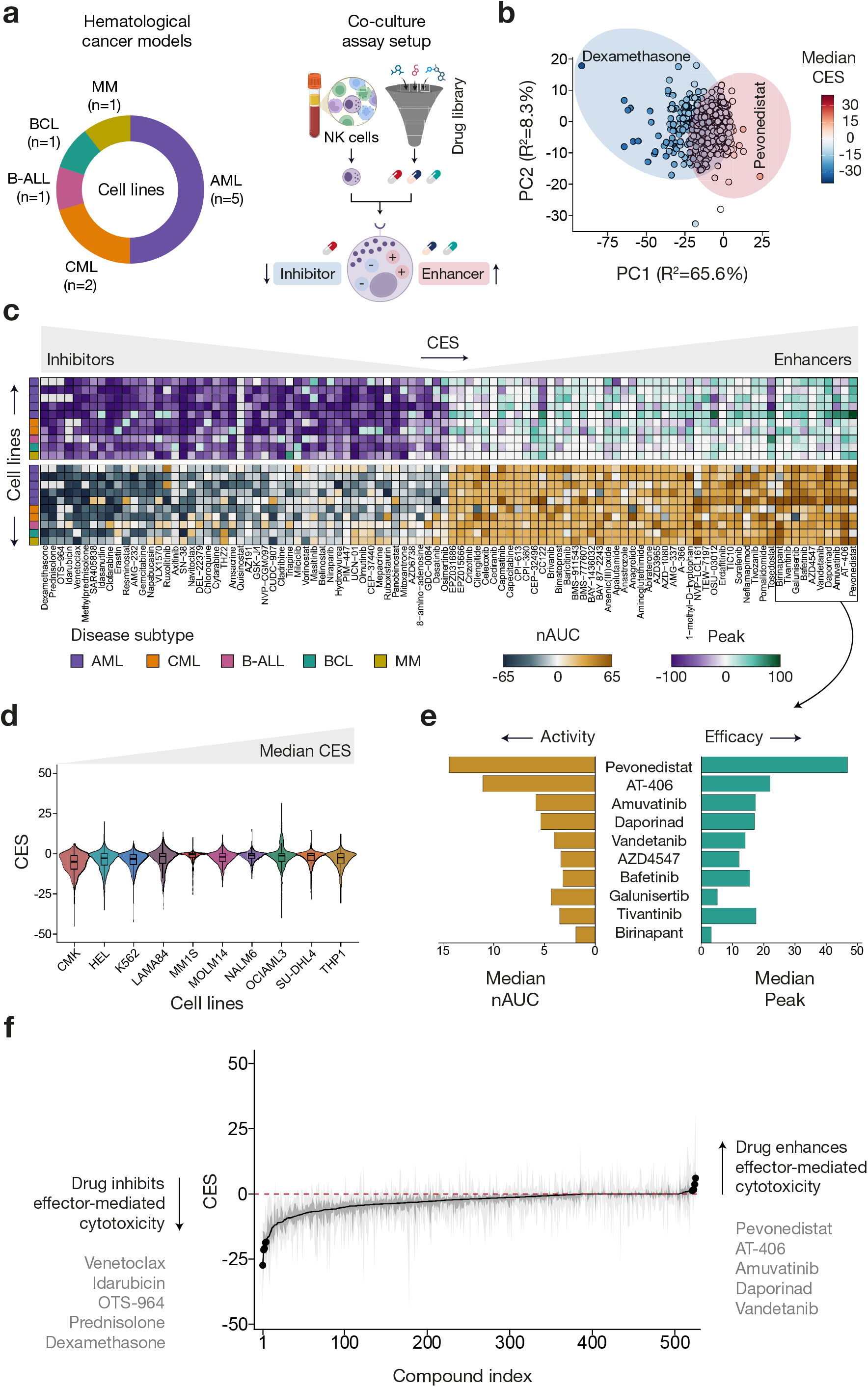
Large-scale quantitative profiling of co-culture drug responses across hematological cancer models. (**a**) Overview of the large-scale co-culture screening framework used for the development of the CES method. Ten hematological cancer cell line models were exposed to a library of 527 compounds under immune effector co-culture conditions with natural killer (NK) cells^31^. Drug responses were quantified across condition-specific activity landscapes and merged into a co-culture screening dataset. (**b**) Principal component analysis (PCA) of median CES values, where each point represents a compound positioned according to its global co-culture efficacy profile. Drugs enhancing NK-cell cytotoxicity (positive CES) are highlighted in red, whereas those inhibiting NK-cell cytotoxicity (negative CES) are shown in blue. (**c**) Maximal effect (Peak) and cumulative activity (nAUC) of the top 50 and bottom 50 compounds ranked by median CES, enabling direct comparison of inhibitory and enhancing profiles. (**d**) Distribution of CES values across all 527 screened compounds for each hematological cancer cell line. Cell lines are ordered by increasing median CES (left to right), illustrating differential susceptibility to modulation of effector-mediated cytotoxicity. Violin plots depict the full distribution of CES values per cell line, with embedded boxplots indicating the median and interquartile range (IQR). (**e**) Cumulative activity and efficacy summaries for ten representative compounds selected from the CES-ranked landscape. (**f**) Global landscape of CES values where compounds are ranked along the x-axis by median CES per compound. Shaded ribbons represent the spread of CES values across the 10 cell lines; the dark gray ribbon denotes the interquartile range (IQR) and the light gray ribbon denotes the full CES range across models. The dashed red line marks CES = 0. Black points show the top five enhancers and bottom five inhibitors, with compound names annotated.

Based on the Gaussian mixture-modeled dose-response profiles, CES values were computed for each compound across all three experimental conditions, enabling systematic comparison of effector-target-drug interactions (**Supplementary Data 3**). Principal component analysis (PCA) of drug-level CES profiles revealed a global activity gradient separating enhancers from inhibitors of selective cytotoxicity, independent of hematopoietic lineage (**Fig. 2b**). Consistent with our previous findings^31^, the NEDD8-activating enzyme (NAE) inhibitor pevonedistat led to consistently high CES values and clustered among other strong enhancers, including SMAC mimetics, whereas glucocorticoids, such as dexamethasone, methylprednisolone, and prednisolone, were among the strongest inhibitors of selective cytotoxicity. The global interaction landscape was biased toward negative CES values, with the CMK co-culture screen displaying the strongest inhibitory profile (mean CES = -6.12; 408 inhibitors) and the single most negative drug response in the screen (selumetinib CES = -45.1), while the MM1S screen showed the least inhibitory effects (mean CES = -1.68). In contrast, the LAMA84 co-culture screen harbored the largest number of 131 enhancer compounds, and the OCIAML3 screen displayed the most extreme enhancement, reaching an average CES of 31.4, indicating pronounced potentiation of effector-mediated cytotoxicity (**Supplementary Table 1**).

To investigate the pharmacological landscape of the blood cancers, we ranked the compounds by their global median CES and visualized the top and bottom 50 drugs across the cell lines (**Fig. 2c**). Visualization of maximal efficacy (Peak) and cumulative activity (nAUC) across the models revealed coherent response patterns while preserving the transition from inhibitory to strongly enhancing interactions. We selected the ten highest-ranking enhancers and decomposed their responses into respective activity and efficacy components (**Fig. 2e**). Among these compounds, nAUC followed a consistent monotonic trend, whereas efficacy varied substantially, indicating heterogeneity in the maximal magnitude of the enhancer effect despite broadly similar cumulative activity profiles.

Among the strongest enhancers, the NAE inhibitor pevonedistat consistently ranked as the top hit, followed by the SMAC mimetic AT-406, the multi-kinase inhibitor amuvatinib, and the NAMPT inhibitor daporinad. Several additional SMAC mimetics appeared prominently among the highest CES-ranked compounds, supporting their strong potentiation of target-selective cytotoxicity, consistent with their underlying effects in effector-mediated cytotoxicity modulation^31,32^. In contrast, the inhibitory end of the landscape was dominated by glucocorticoids, including dexamethasone, prednisolone, and methylprednisolone, alongside targeted and cytotoxic agents such as OTS-964, idarubicin, and venetoclax, suggesting suppression of NK cell-mediated killing in co-culture conditions (**Fig. 2f, Supplementary Fig. 2b, Supplementary Data 4**). Examination of the CES response profiles together with the corresponding CES-derived effective dose profiles provided an additional quantitative layer, enabling direct identification of the most beneficial dosage for each compound across individual cell models (**Supplementary Fig. 2c**).

We next examined the distribution of CES values across all screened compounds to assess model-specific sensitivity to effector-mediated modulation (**Fig. 2d**). The resulting distributions highlighted a broad spectrum of interaction landscapes, demonstrating that while the majority of compounds exert neutral or mild inhibitory effects, a distinct subset of drugs drives strong phenotypic enhancement or inhibition of target-selective cytotoxicity (**Fig. 2f**). Pathway-level analysis of the top enhancer and inhibitor compounds per cell model revealed recurrent pharmacological classes underlying both extremes of the landscape. Enhancer responses were dominated by NAE inhibitors, NAMPT inhibitors, and SMAC mimetics, consistent with recent findings establishing NEDD8 pathway inhibition as an enhancer of NK-cell cytotoxicity across hematological malignancies^31^, whereas inhibitory responses were enriched for glucocorticoids and MDM2 inhibitors, with one to four compounds per class identified in each cell line (**Supplementary Fig. 2d, Supplementary Table 2**). Despite these shared pharmacological trends, CES profiles showed moderate concordance across the five AML models (Spearman ρ = 0.35–0.57; **Supplementary Fig. 2e**), while the two CML models showed weak correlation (Spearman ρ = 0.27; P < 0.0001; **Supplementary Fig. 2f**), highlighting substantial cell context-specific heterogeneity in effector-drug interactions.

### CES is reproducible across screening resolutions and effector-to-target ratios

To evaluate the robustness of CES, we applied the method to an independent validation dataset derived from our previously reported co-culture screen^31^ (**Fig. 3a**). In this experiment, a panel of 36 compounds was newly profiled across the same ten hematological cancer cell lines using a higher dose resolution to assess whether CES-derived interaction profiles and drug rankings remain consistent.

**Figure 3.**
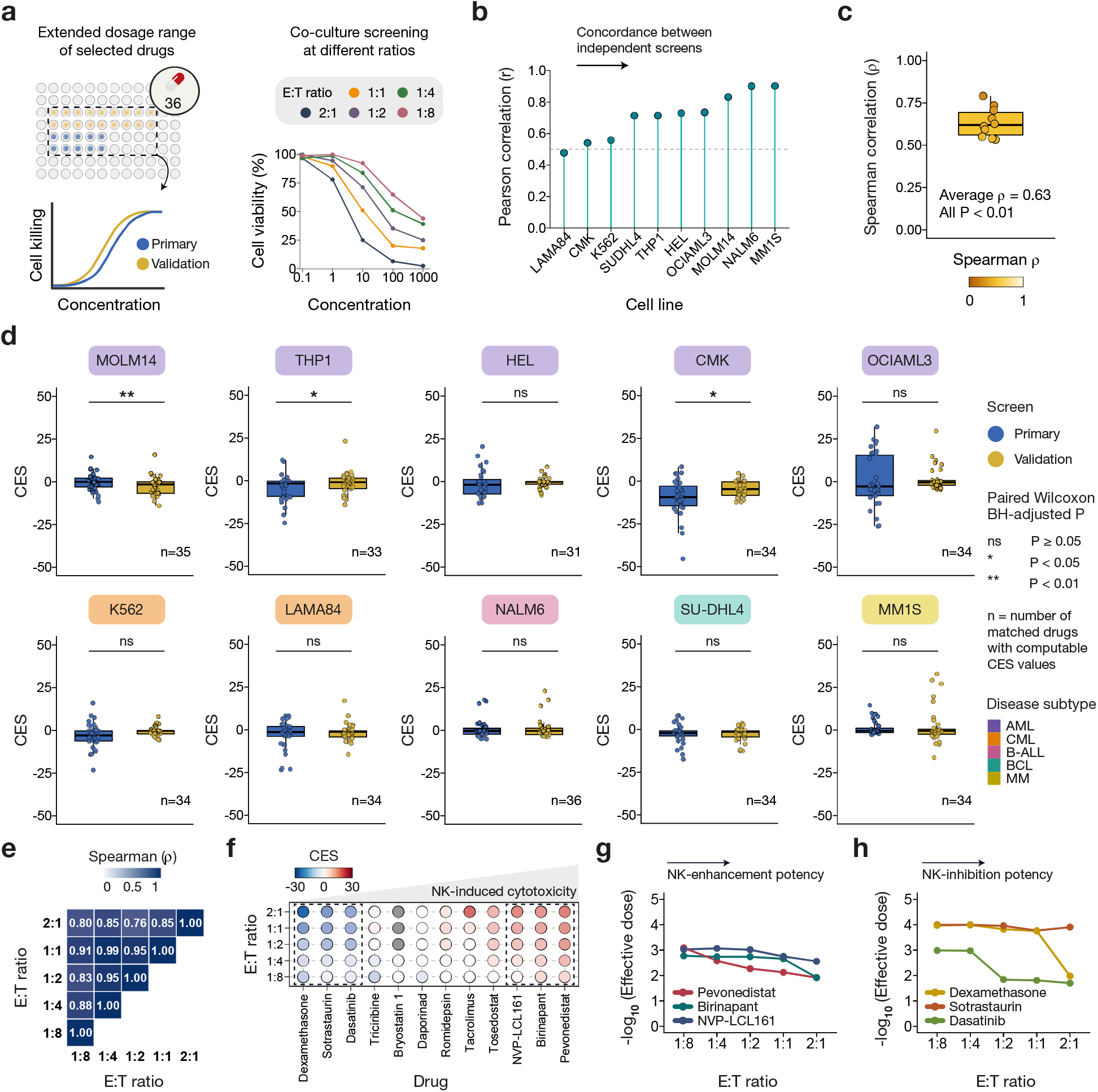
Robustness of the scoring method across independent screens and effector-target conditions. (**a**) Schematic overview of the validation strategy used to assess therapeutic CES robustness. Left: the primary co-culture screen was re-evaluated in an independent validation experiment using a pre-selected panel of 36 compounds profiled at higher dose resolution (nine concentrations) across the hematological cancer cell line panel. Right: CES was additionally computed across varying E:T ratios in the OCIAML3 model to evaluate the stability of immune-dependent drug effects under varying immune pressure. (**b**) Pearson correlation coefficient (*r*) for the 36 tested compounds between the primary and validation datasets, calculated independently for each of the 10 cell lines. The horizontal dashed gray line indicates the correlation threshold of *r* = 0.5. (**c**) Spearman correlation coefficients (ρ) quantifying concordance of drug-level CES values between the primary and validation screens across hematological cancer cell line models. CES values were matched by drug within each model and correlations were computed across paired measurements (31–36 compounds per model). Each point represents one cell line; the boxplot summarizes the distribution of correlation coefficients across models. (**d**) Comparison of CES values between the primary and validation screens. Each point represents one matched compound yielding finite CES values in both datasets (31–36 compounds per model). Y-axis limits were fixed to [−50, 50] to facilitate direct visual comparison across all cell lines. Statistical differences in score distributions were evaluated within each model using a paired Wilcoxon signed-rank test, with P values adjusted for the ten comparisons using the Benjamini–Hochberg (BH) procedure. (**e**) Spearman correlation matrix of CES values across five E:T ratios in the OCIAML3 model across the 36 tested compounds using pairwise complete observations. All pairwise correlations were statistically significant (*P* < 0.05). (**f**) CES values for 12 selected compounds profiled across five E:T ratios in the OCIAML3 model. Drugs are ordered by average CES across conditions, and the E:T ratios are arranged from low to high effector pressure. (**g**) Effective dose (nM) of representative enhancer compounds across E:T ratios in the OCIAML3 model. (**h**) Effective dose (nM) of representative inhibitor compounds across E:T ratios in the OCIAML3 model.

To quantify agreement between the primary and validation screens, we applied CES to the validation screen and compared 36 matched compounds across the ten cell lines (**Fig. 3b-d, Supplementary Data 5**). Pairwise analysis of these matched profiles revealed robust agreement, yielding positive Pearson correlation coefficients across all cell models (*r* = 0.48– 0.90), with most cell lines exhibiting strong reproducibility (**Fig. 3b**) and overall distribution stability (**Fig. 3d**). Significant shifts in CES distributions were observed in only three of the ten models (MOLM14, THP1, and CMK; BH-adjusted P < 0.05), whereas the remaining seven showed no significant deviation. Spearman rank correlations confirmed strong preservation of relative compound efficacy ranking between the primary and validation datasets (ρ = 0.53–0.79, all P < 0.01; **Fig. 3c**), indicating robust performance of the scoring framework at higher dose resolution. Consistent with this finding, direct comparison of raw inhibition measurements confirmed statistically indistinguishable response distributions between the primary and validation screens across both co-culture and target monoculture settings (**Supplementary Fig. 3a**,**b**).

We next evaluated CES stability under varying immune pressure by re-profiling selected compounds across multiple effector:target (E:T) ratios in the OCIAML3 cells (**Fig. 3e,f**). Overall, CES rankings remained largely consistent across conditions, although interaction magnitudes decreased at lower effector pressure, where target cells started to dominate the co-cultures (**Supplementary Data 6**). For example, the top enhancer pevonedistat exhibited a decrease in CES from 27 to 8, and a similar trend was observed across other top hits. Inhibitor compounds showed comparable attenuation under reduced effector pressure, with dexamethasone shifting from a CES of -45.7 to -13.6. Despite these magnitude changes, the cross-condition concordance of drug responses remained high, with Spearman correlations ranging from 0.76 to 0.99 across the E:T ratios (**Fig. 3e**). Consistently, hierarchical clustering of peak response and nAUC profiles revealed high similarity across the E:T ratios (**Supplementary Fig. 3c, d**). This attenuation is consistent with reduced effector-dependent dynamic range at lower E:T ratios. These results show that CES preserved the relative ranking of compounds, while retaining the context-dependent magnitude changes, rather than normalizing them away.

Analysis of CES-derived effective doses further supported the expected immune-dependent effects. Top-enhancer compounds exhibited a marked increase in the required effective dose as the effector pressure decreased **(Fig. 3g**). For example, pevonedistat achieved maximal enhancement at an effective dose of approximately 83 nM at an E:T ratio of 2:1, increasing to roughly 1,235 nM at 1:8, reflecting a reduced pharmacological leverage available under a limited number of effector cells. Antagonistic compounds displayed a similar trend, requiring substantially higher concentrations to achieve maximal inhibition of target-selective cytotoxicity as effector pressure was reduced (**Fig. 3h**). For instance, the effective dose of dexamethasone increased from approximately 96 nM at a 2:1 ratio to 9,304 nM at 1:8, while dasatinib shifted from roughly 50 nM to 965 nM across the same conditions. Together, these results demonstrate that CES preserves consistent drug-response rankings across varying dose resolutions and effector-to-target ratios, while quantitatively capturing the physiological shifts in pharmacological potency that occur as the effector pressure varies.

### Application of CES to immune co-culture and antiviral host-pathogen screens

To evaluate the generalizability of CES across various biological systems, we next analyzed two external screening datasets representing distinct multicellular and host–pathogen contexts (**Fig. 4a**). The first screen investigated pharmacological modulation of CD19 CAR T-cell-mediated killing in NALM6 leukemia cells^28^. The second dataset measured host-cell viability rescue in SARS-CoV-2-infected Vero E6 cells^20^. Because these two studies lacked effector-only control conditions, CES was applied in its two-condition mode. As a result, these analyses quantify differential co-culture or infection-associated activity relative to the matched reference condition, yet cannot resolve direct effector-cell toxicity.

**Figure 4.**
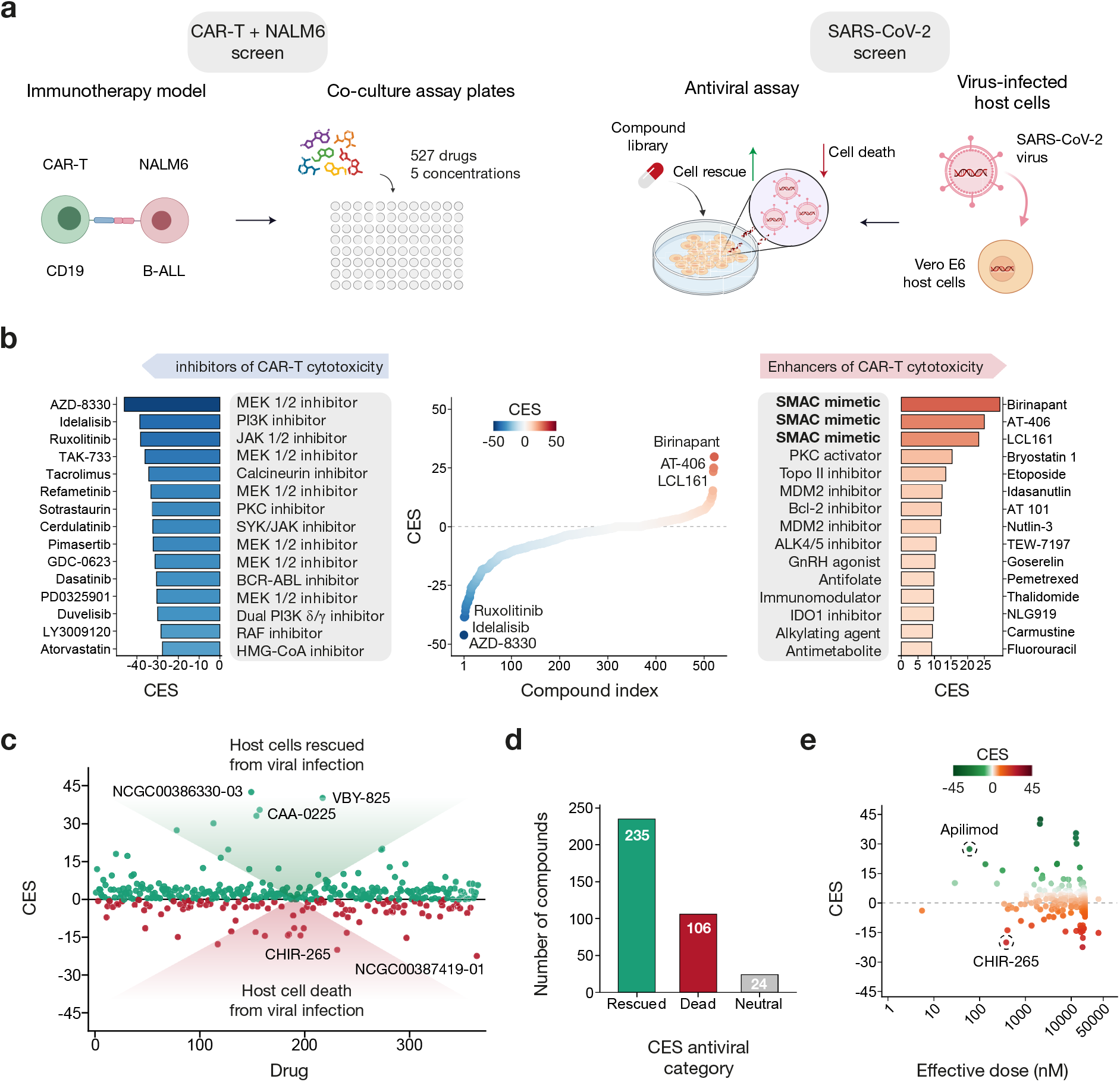
Application of CES to independent immune co-culture and antiviral screening datasets. (**a**) CES was applied to two external two-condition datasets. Left: CD19 CAR T-cell co-culture drug screen in NALM6 leukemia cells, in which target cells were cultured alone or together with anti-CD19 CAR T cells and target-cell viability was measured after 24 h using a luciferase-based readout. Right: SARS-CoV-2 host-cell rescue screen in Vero E6 cells, in which CES quantified differential host-cell viability under infected and reference conditions. Because neither dataset includes an effector-only condition, CES was computed in its two-condition mode (see **Methods**). (**b**) Ranked CES landscape across compounds in the CAR T-cell co-culture dataset. Compounds are ordered by CES rank, with the left and right panels highlighting the top 15 inhibitors and top 15 enhancers of CAR T-cell-mediated cytotoxicity, respectively. (**c**) Distribution of CES values across compounds in the antiviral screening dataset. Each point represents one PubChem-indexed compound. Positive values indicate host-cell rescue from viral infection, whereas negative values indicate dominant viral cytopathic effects. The horizontal reference line denotes CES = 0, indicating neutral activity. (**d**) Bar plot summarizing CES-derived antiviral response categories across the screened compounds. Bars represent the number of hits classified as Rescued (CES > 0), Dead (CES < 0), or Neutral (CES = 0). (**e**) Relationship between CES and effective dose in the antiviral screening dataset.

Application of CES to the CAR T-cell co-culture dataset revealed a clear pharmacological landscape separating compounds that either enhance or inhibit CAR T-cell-mediated cytotoxicity (**Fig. 4b, Supplementary Data 7**). Several SMAC mimetics ranked among the strongest positive-scoring compounds (CES = 23-30), together with bryostatin 1 and multiple MDM2-class inhibitors, consistent with drug classes previously implicated in modulation of CAR T-cell activity^28^ and in KRAS-mutated patient-derived organoids (PDOs)^42^. Strikingly, the negative end of the ranking was dominated by MEK1/2 inhibitors: six compounds from this class clustered among the top 15 inhibitors with markedly negative scores (mean CES = -35.4), consistent with the role of MAPK/ERK signaling in T-cell effector function^43^. AZD-8330 emerged as the most potent inhibitor, with an effective dose of approximately 17 nM. The modeled co-culture dose-response curves for the top four enhancers and inhibitors illustrated the dose-dependent magnitude of this immunomodulation (**Supplementary Fig. 4a, b**). Relating CES components to pharmacological potency further illustrated these effects: the top enhancer birinapant had an effective dose of approximately 342 nM, whereas bryostatin 1 showed the lowest effective dose among the top hits (∼40 nM) (**Supplementary Fig. 4c, d**). Comparison of efficacy and activity across compounds revealed strong correlation (Spearman ρ = 0.915, *P* < 0.0001; **Supplementary Fig. 4e**), indicating that CES-based metrics capture complementary yet partly redundant dimensions of drug response in the co-culture system.

We next applied CES to an antiviral screening dataset in which compounds were evaluated for their ability to rescue viability of SARS-CoV-2-infected Vero E6 host cells (**Supplementary Data 8**). The CES landscape separated compounds with positive host-cell rescue profiles from compounds with negative or neutral infection-associated response profiles (**Fig. 4c**). Notably, the highest-ranking compounds exhibited substantially larger positive CES values than the most negative scores observed among non-rescuing drugs, highlighting a strong asymmetry in the magnitude of rescue versus cytotoxic responses. Quantifying this polarity across the screened compounds further emphasized the predominance of antiviral rescue events: 236 compounds displayed positive CES, indicating protection of the host cell; a total of 105 compounds showed negative CES, reflecting dominant viral cytopathic effects; and 24 compounds exhibited neutral responses (CES = 0) (**Fig. 4d**).

Relating CES values to model-estimated effective concentrations showed that many of the highest-ranked rescue compounds were active only at higher tested concentrations. Most top rescue hits exhibited effective doses between approximately 2,055 and 12,500 nM, with the notable exception of apilimod, which showed markedly higher potency at 60 nM) (**Fig. 4e**). A similar pattern was observed among compounds associated with dominant cytopathic outcomes, where effective doses were also typically high, except for CHIR-265 (∼381 nM). Examination of the corresponding antiviral dose-response profiles of host cells further illustrated various response dynamics: the four highest-ranked rescue compounds demonstrated near-complete restoration of host-cell viability at intermediate drug concentrations, indicating effective suppression of virus-induced cytopathic effects (**Supplementary Fig. 4f**). In contrast, the four compounds with the strongest negative CES showed no detectable rescue within the tested dose range, with host-cell viability remaining near zero under the infection conditions (**Supplementary Fig. 4g**). Taken together, CES resolves both antiviral rescue and cytopathic response profiles, extending its applicability beyond immune co-culture screens to host-pathogen drug discovery contexts.

### CES improves compound coverage and specificity relative to conventional pharmacological metrics

To place CES within a broader drug sensitivity scoring landscape, we re-analyzed a recently published high-throughput screen of 527 oncological compounds co-cultured with primary NK cells and K562 chronic myeloid leukemia (CML) cells^32^. We first sought to evaluate the complementary roles of the two CES modes, therapeutic CES (used in the previous sections) and mechanistic CES (see **Methods**), prior to benchmarking against conventional metrics (**Fig. 5a**). This comparison was evaluated in terms of distinguishing targeted immunomodulation from general cytotoxicity, defined as substantial direct toxicity against the effector NK cells (DSS ≥ 10; see **Methods**). Because conventional metrics capture distinct dimensions of drug activity, ranging from absolute potency to differential toxicity across cellular contexts, we systematically benchmarked our dual CES modes against these established pharmacological measures.

**Figure 5.**
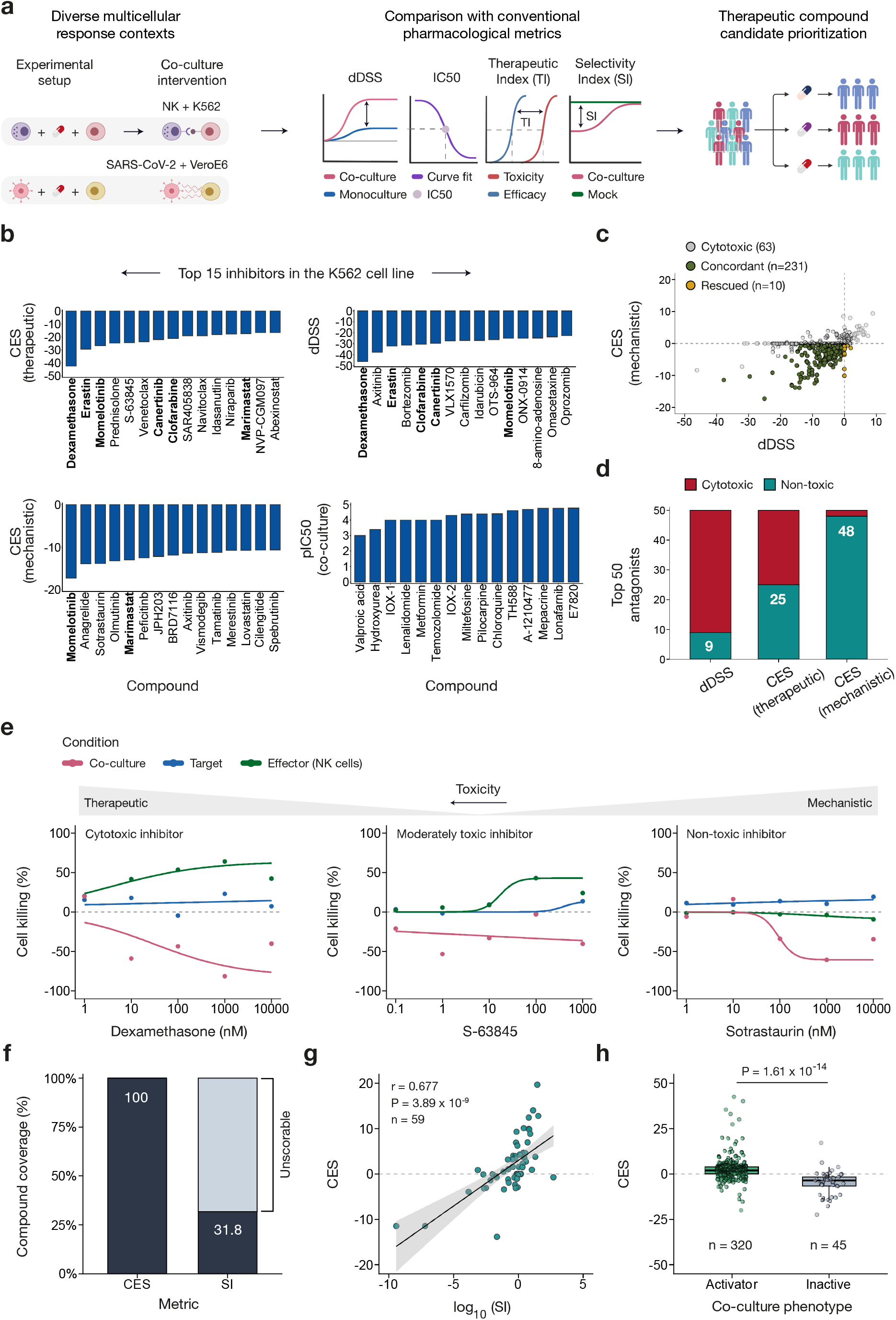
CES resolves therapeutic selectivity and drug response mechanisms across immune and antiviral contexts. (**a**) Schematic overview of the CES framework and its comparison with conventional pharmacological metrics across diverse multicellular response contexts. CES quantifies context-specific drug responses from co-culture dose-response profiles and enables direct comparison with established measures, including the half-maximal inhibitory concentration (IC_50_), differential drug sensitivity score (dDSS), therapeutic index (TI), and antiviral selectivity index (SI). (**b**) Top 15 inhibitors in the K562 NK-cell co-culture screen (among 527 compounds) ranked by therapeutic CES, mechanistic CES, dDSS, and co-culture pIC_50_ (-log_10_ of the IC_50_ in molar units). pIC_50_ is included as the conventional single-curve baseline, whereas therapeutic CES, mechanistic CES, and dDSS integrate information across full dose-response profiles and reference conditions. Compounds shared between therapeutic CES, and the alternative scoring metrics are emphasized with bold text. (**c**) Identification of non-toxic antagonists of effector-mediated cytotoxicity in the K562 screen (517 compounds with complete data). Scatterplot of mechanistic CES versus dDSS across all compounds. Cytotoxic compounds (defined as DSS ≥ 10 against the NK-cell control condition) are shown in gray (63 compounds), concordant non-toxic antagonists in olive (231 compounds), and non-toxic antagonists flagged by mechanistic CES (CES < 0) but scored as neutral or enhancing by dDSS (dDSS ≥ 0) in gold (10 compounds). Dashed lines indicate the zero threshold on each axis. (**d**) Comparison of toxicity profiles among the top 50 predicted antagonists identified by dDSS, therapeutic CES, and mechanistic CES in the K562 screen. The stacked bar chart displays the number of broadly cytotoxic compounds (DSS ≥ 10 of the effector cells; red) versus non-toxic antagonists (teal) prioritized by each metric. (**e**) Dose-response curves of dexamethasone, S-63845, and sotrastaurin in the K562 co-culture screen. Three conditions are shown per example compound, illustrating distinct antagonist archetypes: dexamethasone exhibits broad cytotoxicity, S-63845 shows mild toxicity, and sotrastaurin selectively inhibits effector-mediated killing in co-culture with no direct cytotoxicity on effector cells. (**f**) Proportion of antiviral compounds with computable IC_50_-based selectivity index (SI) compared with CES coverage. SI was calculable for 116 of 365 compounds (31.8%), whereas CES produced quantitative scores for all compounds. (**g**) Association between CES and log_10_(SI) for the subset of 59 compounds yielding strictly positive SI values. The Spearman correlation coefficient (ρ) is reported. (**h**) Distribution of CES values stratified by antiviral phenotype (Activator vs Inactive). Differences were tested with a two-sided Wilcoxon rank-sum test.

When examining compounds that enhance NK-cell-mediated killing, the top-15 enhancers showed a substantial overlap across therapeutic CES, mechanistic CES, and dDSS, albeit with differences in their ranking (therapeutic vs dDSS: 12 shared; therapeutic vs mechanistic: 11 shared; Fisher’s exact test, *P* < 10−^16^; **Supplementary Fig. 5a**). Notably, the SMAC mimetics birinapant and NVP-LCL161 were consistently recovered across all three metrics, consistent with their role as potent enhancers of NK-cell cytotoxicity^28,32^ (**Supplementary Fig. 5d**). In contrast, the top 15 inhibitors revealed striking divergence across the metrics (**Supplementary Data 9**; **Fig. 5b**). Therapeutic CES shared five compounds with dDSS (Fisher’s exact test, *P* = 2.3 × 10^−5^) and only two with mechanistic CES (P = 0.064), while no compounds overlapped with the co-culture pIC_50_ ranking. The dDSS-recovered inhibitors were heavily enriched for broadly cytotoxic agents, such as proteasome inhibitors and anthracyclines, whereas therapeutic CES uniquely identified targeted, non-cytotoxic agents, including BCL2/MCL1, MDM2, and PARP inhibitors. Mechanistic CES covered an entirely distinct profile dominated by kinase inhibitors that modulate NK-cell signaling without direct cytotoxicity, such as the PKC inhibitor sotrastaurin. Furthermore, therapeutic CES showed a modest positive correlation with the co-culture therapeutic index (TI), defined as the ratio of NK-cell IC_50_ to co-culture IC_50_ (Spearman *ρ* = 0.142, *P* = 0.0012; **Supplementary Fig. 5b**), and stronger correlation with TI defined against the target-cell monoculture IC_50_ (*ρ* = 0.30, *P* = 2.96 × 10^−12^; **Supplementary Fig. 5c**). Compounds in the highest therapeutic CES quartile exhibited significantly greater TI values than those in the lowest quartile (Wilcoxon rank-sum test, *P* = 4.2 × 10^−3^; **Supplementary Fig. 5e**). Together, these results demonstrate that CES effectively integrates the full dose-response landscape against condition-matched controls, rather than summarizing selectivity through a single IC_50_ ratio.

We next evaluated the accuracy of identifying non-toxic antagonists of effector-mediated cytotoxicity (**Fig. 5c, d**). Among non-toxic compounds flagged as antagonists by mechanistic CES (CES < 0), 10 compounds (4.1%) were scored as neutral or enhancing by dDSS (dDSS ≥ 0), while the remaining 231 (95.9%) were scored as antagonists by both metrics (**Fig. 5c**). Analysis of the top-50 predicted antagonists per metric revealed a striking resolution gradient (**Fig. 5d**), where the dDSS top-50 was dominated by broadly cytotoxic compounds (41 of 50; 82%), therapeutic CES yielded an even split (25 cytotoxic, 25 non-toxic), and mechanistic CES nearly exclusively isolated genuine, non-toxic antagonists (48 of 50; 96%). This selectivity was robust across ranking cutoffs with mechanistic CES, identifying on average only 2.6% cytotoxic compounds across the top 100 antagonists, compared with 53.4% for therapeutic CES and 75.7% for dDSS (**Supplementary Fig. 5g, h**). Dose-response curves of the representative antagonists clearly illustrate the mechanistic basis for this divergence (**Fig. 5e**). Dexamethasone exhibited a high toxicity to NK cells and a reduced target-cell killing in co-cultures, representing a toxic response rather than true immunomodulation. S-63845 displayed modest effector toxicity and a slight reduction in co-culture killing. In contrast, sotrastaurin showed no toxicity in either target or effector monocultures, yet it reduced target-cell killing in the co-culture, allowing target cells to proliferate. This genuine immunomodulatory pattern was captured exclusively by the mechanistic CES. These results establish mechanistic CES as the preferred metric for identifying genuine antagonists of effector-mediated cytotoxicity, effectively preventing the misclassification of broad chemotherapeutics as specific immunomodulators.

When extending this evaluation to an antiviral screening dataset of 365 compounds tested in virus-infected host cells, we observed similar advantages of CES over conventional metrics. The classical selectivity index (SI) could only be computed for 116 of 365 compounds (31.8%), due to missing or undefined IC_50_ values, whereas CES generated quantitative scores for all compounds (**Fig. 5f**). Of the 116 compounds, 59 yielded positive values suitable for log_10_-transformation. Across these, CES was positively correlated with antiviral selectivity (Spearman *ρ* = 0.677, *P* = 3.89 × 10^−9^; **Fig. 5g**). Compounds classified as antiviral activators displayed significantly higher CES values than inactive compounds (median 1.95 vs -3.61; Wilcoxon rank-sum test, *P* = 1.6 × 10^−14^; **Fig. 5h**), and compounds exhibiting host-cell cytotoxicity in the control condition showed reduced CES values relative to non-cytotoxic compounds (median CES 0.00 vs 2.45; Wilcoxon rank-sum test, *P* = 1.8 × 10^−9^; **Supplementary Fig. 5f**). Together, these analyses extend the applicability of CES to host-pathogen screening contexts, where conventional selectivity metrics are limited.

## Discussion

We developed the Co-culture Efficacy Score (CES), a model-based metric and analytical platform designed to quantitatively capture complex compound perturbation landscapes across diverse multicellular contexts. By bridging the gap between monoculture and co-culture pharmacological responses, CES provides a generalizable framework for identifying immune-modulating agents for cancer immunotherapy and host-protective compounds for antiviral discovery. Our analysis of co-culture dose-response profiles (**Supplementary Fig. 1, Supplementary Data 1**) revealed highly heterogeneous interaction patterns, where drug treatments can either potentiate or suppress immune-mediated target-cell killing, while simultaneously exerting independent effects on both immune effector and target cancer cells, creating complex pharmacological landscapes that monoculture assays fail to capture. Furthermore, because drug-induced toxicity toward effector cells often confounds co-culture screening, CES incorporates an explicit quantitative penalty for compounds that diminish apparent co-culture efficacy by impairing effector-cell viability. To ensure this methodology is immediately deployable, the CES framework is implemented as an open-source computational tool alongside a standalone, interactive web application (https://ces.aittokallio.group). By automating the analysis of co-culture datasets, the CES web-application lowers technical barriers and empowers the broader research community to discover combinations of drug and cell-based therapies.

Our application of therapeutic CES to an NK-cell co-culture screening dataset^31^ revealed extensive heterogeneity in effector-drug interactions, consistent with the broad pharmacogenomic variability observed in cancer cell models^44,45^. While most compounds predominantly inhibited NK-mediated cytotoxicity (**Fig. 2d, Supplementary Table 1**), CES successfully prioritized shared mechanistic pathways across cancer models and pinpointed model-specific vulnerabilities (**Fig. 2f**; **Supplementary Fig. 2d**). For example, pevonedistat (a NEDD8-activating enzyme inhibitor) and a broader set of SMAC mimetics consistently emerged as the top-ranked enhancers across nearly all hematological models. Beyond these shared enhancers, CES identified distinct disease-specific patterns, such as NAE and NAMPT inhibitors in AML; SMAC mimetics in CML; and MET, EGFR, and AKT inhibitors in B-cell lymphoma. Notably, OCIAML3 cell line stood out for the breadth of its enhancer landscape, spanning a highly diverse set of pathway classes. In contrast, the inhibitory end of the landscape was broadly convergent across the models, with glucocorticoids and MDM2 inhibitors emerging as the dominant classes in nearly every cell line (**Supplementary Table 2**).

We systematically evaluated the CES outcomes in independent datasets to assess robustness and generalizability. We compared CES values between the primary and validation screens to confirm consistent drug interaction profiles across hematological cancer models (**Fig. 3b-d, Supplementary Fig. 3a**,**b**). Although interaction magnitudes decreased at lower effector-to-target ratios, as expected, compound rankings were largely preserved (**Fig. 3f**). Importantly, CES quantitatively captured the physiological shifts in drug potency, with effective concentrations required for peak activity increasing as baseline immune leverage diminished. Methodologically, these shifts underscore the necessity of calibrating assays to an intermediate baseline of effector cytotoxicity. This baseline provides the dynamic range to confidently identify both strong enhancers and potent inhibitors, without requiring the use of non-physiological drug concentrations. Even under conditions with reduced effector activity, CES retained the ability to robustly identify selective hits, highlighting its potential for reliable compound prioritization. Application to the CAR T-cell co-culture dataset revealed a high degree of concordance with previously established enhancers, while effectively prioritizing compound efficacy and distinguishing top inhibitors. Notably, we identified the MEK1/2 drug class as strong antagonists of CAR T-cell activity in NALM6 cells (**Fig. 4b**; **Supplementary Data 7**). Application of the metric to a host-pathogen antiviral screening dataset^20^ further illustrated its performance beyond cancer immunology (**Fig. 4c-e**). We successfully differentiated compounds that restore host-cell viability in infected cells from those associated with viral cytopathicity, a critical distinction in antiviral drug discovery, where the suppression of viral replication must be balanced with the preservation of host cellular viability^46-48^.

Quantitative dose-response analysis of drug responses in cell-based screens has traditionally relied on potency metrics, most commonly IC_50_ concentration^49^. However, multiparametric analyses of dose-response behavior have shown that additional features, such as slope of the response curve (Hill coefficient) and maximal effect (Emax), capture systematic variation related to drug class and cellular context that is not reflected in IC_50_ alone^50-52^. Summary metrics integrating these parameters, such as area under the inhibitory curve (AUC) and drug sensitivity score (DSS)^35^, have therefore become widely used in large-scale pharmacological profiling studies and construction of pharmacogenomic resources^40^. Recent efforts to improve quantitative drug-response analysis address variability in cell growth dynamics and experimental noise to enhance the robustness and comparability of screening measurements^53,54^. Nevertheless, these approaches were primarily developed for monoculture assays and therefore do not account for effector-mediated cytotoxicity modulation that arises when multiple cell populations contribute to the measured response. These limitations motivated the development of CES, a model-based scoring framework designed to quantify drug sensitivity in co-culture systems, where pharmacological effects emerge from the interplay between drug action, effector-mediated cytotoxicity, and tumor-cell sensitivity.

A central challenge in analyzing co-culture drug screens is that conventional monoculture-derived metrics often misclassify non-specific cytotoxic compounds as immune antagonists. Agents that broadly impair cell viability reduce effector-cell fitness, thereby producing a spurious signature of immunomodulation. Our comparative analyses demonstrated that this confounding signal severely distorts traditional area-based scores. Among the top 50 antagonists ranked by each metric, only 9 of the dDSS hits represented true non-toxic antagonists, whereas therapeutic and mechanistic CES identified 25 and 48 antagonists, respectively (**Fig. 5d**). Mechanistic CES resolved these confounding effects almost perfectly by adjusting for effector-only responses, rather than simply penalizing effector toxicity; and it additionally recovered 10 genuine antagonists that dDSS misclassified as neutral or enhancing (**Fig. 5c**). These results establish mechanistic CES as the preferred metric for identifying antagonists of effector-mediated cytotoxicity in co-culture drug screens, successfully distinguishing true pharmacological antagonism from non-specific cytotoxicity. However, therapeutic CES serves as the broadly applicable version that directly penalizes effector-cell toxicity, making it suitable for prioritizing compounds with a favorable therapeutic window in large-scale co-culture screens.

A second limitation of conventional metrics is their dependence on absolute IC_50_ estimates. Although relative IC_50_ values can be computed even when dose-response curves fail to reach the 50% inhibition threshold, these estimates become highly unstable and unreliable for compounds with low maximal efficacy^46,55^. In the antiviral dataset, we showed how the classical selectivity index (SI) could only be computed for 116 of 365 compounds (31.8%), excluding the majority of the screened compounds from selectivity-based prioritization (**Fig. 5f**). In contrast, CES does not require IC_50_ estimates, and generated quantitative scores for all 365 compounds. Importantly, within the 116-drug subset where SI could be calculated, CES strongly correlated with the traditional selectivity rankings (Spearman ρ = 0.677; **Fig. 5g**), confirming that CES accurately captures established pharmacological selectivity, while vastly expanding compound coverage. These two gaps, misclassification of non-specific cytotoxicity and exclusion of compounds with undefined curve parameters, reflect distinct weaknesses of conventional pharmacological metrics in multicellular screening contexts. In contrast, CES enables robust compound identification in co-culture settings, where target and effector responses are intrinsically coupled within a single multicellular assay, addressing both gaps within a single scoring framework.

Functional screening technologies are rapidly evolving from simplified monoculture assays toward more advanced experimental platforms that better capture immune-mediated responses between multiple cell populations. Co-culture systems in cancer immunology enable systematic interrogation of how pharmacological perturbations reshape cellular interactions and influence immune-mediated cytotoxicity. While the current CES framework provides a robust analytical foundation for existing co-culture systems, next-generation tumor models are increasingly incorporating additional microenvironmental components such as stromal cells and regulatory immune populations. Capturing these multidimensional interactions within organoid models will require further development of scoring approaches to address the growing biological complexity. As functional screening continues to scale toward higher-throughput formats and diverse patient-derived models, the immediate demand for robust and accessible analytical tools remains critical. For these existing and emerging applications, the CES methodology and its web application offer an immediately scalable and automated solution. By automating the rigorous modeling required by these experiments, CES framework enables researchers to efficiently resolve complex drug-effector-target interactions across diverse co-culture assay environments.

## Materials and Methods

### Co-culture screening data for the development of the CES model

The experimental co-culture datasets utilized to develop and validate the computational framework were originally generated and detailed in a companion biological study^31^. To provide necessary context for the analytical methods, the biological screening procedures are summarized below.

### Co-culture drug sensitivity and resistance testing (DSRT)

High-throughput drug sensitivity and resistance testing (DSRT) was performed at the Institute for Molecular Medicine Finland (FIMM) using three distinct plate formats: (i) co-culture plates containing both target and effector cells, (ii) target monoculture plates, and (iii) effector-only toxicity plates. A panel of 527 FDA-approved and investigational oncological compounds (FO5A FIMM oncology library), spanning diverse mechanistic classes, was screened across five concentrations. These were distributed on a log_10_-scaled range to enable the evaluation of dose-dependent responses across multiple orders of magnitude. Compounds were pre-dispensed into 384-well plates using acoustic liquid-handling technology. For co-culture experiments, luciferase-expressing target cells were plated either alone or with effector cells at predefined effector:target (E:T) ratios. Monoculture conditions were included to independently quantify drug effects on both target and effector cell-viability. Following 24 hours of incubation at 37 °C, target cell viability was quantified via a luminescence-based readout using ONE-Glo (Promega) on a PHERAstar microplate reader (BMG Labtech). Effector-cell viability was assessed separately using CellTiter-Glo (Promega) under matched exposure conditions.

### Cancer cell line models

Ten hematological cancer cell line models were included in the co-culture DSRT screen: five acute myeloid leukemia (AML) models (MOLM-14, THP-1, HEL, CMK, and OCI-AML3), two chronic myeloid leukemia (CML) models (K562 and LAMA-84), one B-cell precursor acute lymphoblastic leukemia (B-ALL; NALM6), one B-cell lymphoma (SU-DHL4), and one multiple myeloma model (MM1S). All cell lines used for DSRT were transduced to express luciferase to facilitate luminescence-based target viability quantification in mixed cultures.

### Data normalization and quality control

Raw luminescence intensities were normalized to plate-specific controls to obtain relative response values expressed as percentages. Benzethonium chloride (BzCl) served as the positive control for maximal cell death, while DMSO-treated wells were used as negative controls. In co-culture plates, negative controls consisted of DMSO plus NK cells to account for baseline effector-mediated killing, whereas monoculture plates used DMSO alone. These normalized measurements served as the standardized inputs for the CES computational pipeline. Plate-level quality control was rigorously assessed using the Z-prime (Z′) and robust Z-prime (robust Z′) factors, as detailed in **Results Section 2** and in **Supplementary Data 2**.

### Optimization of baseline effector-to-target ratios

Prior to the primary drug sensitivity and resistance testing (DSRT) screens, we determined the optimal effector-to-target (E:T) ratio for each individual target cell line. The objective was to establish a baseline E:T density that yielded approximately 50% inhibition of target cell viability in the absence of drug treatment. Securing this mid-point baseline ensures an optimal dynamic range for the subsequent screens, allowing for the robust detection of both pharmacological enhancers and inhibitors of effector-mediated cytotoxicity. Full experimental procedures are described in the companion study^31^.

### Validation in higher-resolution co-culture screening

To validate findings from the primary high-throughput DSRT screen, a focused panel of 36 compounds was selected from the original 527 compound library and re-screened at nine concentrations. As detailed in the companion biological study^31^, this secondary screen enabled higher-resolution dose-response characterization. Compound selection prioritized agents exhibiting strong differential sensitivity across the hematological panel while excluding those with high intrinsic effector-cell toxicity. Validation experiments were conducted using primary NK cells derived from three independent healthy donors to assess reproducibility across distinct effector sources. Co-culture experiments utilized the optimized E:T ratios established in the primary screen. The increased dose resolution of this validation panel enabled detailed evaluation of response curve morphology and confirmed the computational stability of CES across independent biological replicates.

### Impact of varying E:T ratios on the metric performance

To evaluate the influence of effector cell density on drug response profiles and to benchmark the robustness of the CES, the OCI-AML3 cell line was screened across a broad range of E:T ratios (2:1, 1:1, 1:2, 1:4, and 1:8) in replicate wells. Raw viability measurements at each density were normalized to their corresponding condition-specific co-culture controls to account for baseline effector-mediated killing. The resulting dose-response profiles were analyzed to quantify the impact of effector density on assay dynamic range, curve morphology, and susceptibility to floor and ceiling effects. Profiles were then subjected to Gaussian mixture fitting and automated CES computation to assess the capacity of the metric to capture and adjust for shifting interaction dynamics across effector loads (**Supplementary Data 6**).

### CAR T-cell co-culture screening dataset

To evaluate the CES framework in an engineered effector context, we analyzed an independent, previously reported CD19-targeting CAR T-cell co-culture screening dataset^28^. In this screen, the CD19-positive B-ALL cell line NALM6 (engineered to express luciferase) was incubated either alone or with CAR T-cells in 384-well plates. The cells were treated with an oncology library comprising 527 compounds (both US Food and Drug Administration/European Medicines Agency-approved anticancer drugs as well as investigational and preclinical compounds) across five concentrations spanning a 10,000-fold range. Following a 24-hour incubation, target cell viability was measured via a luminescence-based assay. These normalized viability readouts were subsequently processed through the CES pipeline to identify antagonists and enhancers of CAR T-cell-mediated cytotoxicity.

### Antiviral screening dataset

We used quantitative high-throughput screening data from two complementary PubChem bioassays (AID 1508606 and AID 1508605) to evaluate compound efficacy in infectious diseases such as COVID-19^20^. The primary dataset (AID 1508606) provides the live SARS-CoV-2 cytopathic effect reduction assay, measuring the capacity of therapeutic agents to rescue Vero E6 cell viability during viral infection. The secondary dataset (AID 1508605) provides the cytotoxicity counterscreen conducted in uninfected Vero E6 cells, which is necessary to distinguish true antiviral protection from baseline compound toxicity. For the primary cytopathic effect reduction assay, Vero E6 cells were inoculated with SARS-CoV-2 (strain USA_WA1/2020) at a multiplicity of infection of 0.002. The virus and cell mixture was dispensed into 384-well plates preloaded with the tested compounds, achieving a final density of 4000 cells per well. Following a 72-hour incubation at 37°C and 5% CO_2_, host-cell viability was quantified by measuring cellular ATP content via a CellTiter-Glo luminescent readout. Luminescence data were normalized using DMSO as a negative control and Calpain inhibitor IV as a positive control. Active antiviral compounds protect the host cells from viral-induced cytopathic effects, resulting in preserved viability and higher luminescence signals. Parallel cytotoxicity counterscreens were performed to assess baseline compound safety in the absence of viral infection. Uninfected Vero E6 cells selected for high ACE2 expression were plated at 4000 cells per well with the test compounds and incubated under identical conditions. Viability was assessed using the same luminescent ATP readout. These uninfected control data were normalized against DMSO negative controls and Hyamine positive cytotoxic controls. This dual-assay format ensures that observed therapeutic effects are driven by antiviral mechanisms rather than general cellular toxicity.

### Drug dose-response curve fitting in co-cultures and monocultures

Dose-response curves for target monocultures, effector monocultures, and co-cultures were fitted using a procedure adapted from the Breeze analysis framework^56^ for co-cultures, featuring custom modifications to ensure robust handling of co-culture perturbation data. Raw data were structured by dose, screening context, and inhibition values, with drug concentrations transformed to a log_10_ scale. Inhibition values were constrained between -100 and 100 to prevent extreme outliers from skewing the fit. To maintain numerical stability, duplicated inhibition values were marginally perturbed while preserving rank order. For conditions exhibiting growth enhancement (negative inhibition) at upper doses, the response direction was temporarily inverted to allow consistent estimation of sigmoidal parameters before being transformed back to the original orientation. Initial parameter estimates were obtained with a four-parameter log-logistic model (slope, lower asymptote, upper asymptote, and IC_50_), and these estimates were then refined by bounded nonlinear least-squares optimization using the NL2SOL (‘port’) algorithm in R. Two starting configurations were evaluated, and the fit with the lower residual error was retained. If the log-logistic initialization failed, a four-parameter logistic formulation was substituted, and if the refinement step failed to converge, a Levenberg– Marquardt fallback was used. Following model fitting, predicted response trajectories were generated on a dense concentration grid spanning the experimental dose range. Standard pharmacological parameters (IC_50_, absolute IC_50_, slope, minimum/maximum response) were extracted and transformed to a common efficacy scale. To ensure the biological validity of prioritized hits, an automated secondary quality control filter was applied to exclude artifactual dose-response profiles, specifically targeting compounds that exhibited globally aberrant negative sensitivities or mathematical discordance between the fitted curve trajectories and the resulting Co-culture Efficacy Score (CES). This complete automated curve-fitting architecture serves as the computational backend for the CES platform, generating the standardized, high-quality profiles required for downstream feature extraction.

### Statistical analysis

All statistical analyses were performed in R (version 4.4.0) unless otherwise stated. Drug-response metrics derived from dose-response modeling, including maximal efficacy (Peak), cumulative activity (normalized area under the response curve, nAUC), effective dose, and the Co-culture Efficacy Score (CES), were analyzed across compounds and experimental conditions. Summary statistics are reported as the median with interquartile range (IQR) or as mean values averaged across cell line models, depending on the analysis. The global structure of CES values across compounds was explored using principal component analysis (PCA), a dimensionality reduction method that summarizes variation in high-dimensional datasets through orthogonal linear components^57^. PCA was performed on the matrix of median CES values per compound across cancer cell line models after centering and scaling of variables. Differences between paired experimental conditions were evaluated using paired Wilcoxon signed-rank tests, whereas comparisons between independent groups were assessed using two-sided Wilcoxon rank-sum tests. When multiple hypothesis tests were performed across models, Benjamini-Hochberg false discovery rate (FDR) correction^58^ was applied. To assess whether the overlap between the top 15 compounds identified by each metric exceeded a random expectation, we performed one-sided Fisher’s exact tests on a 2 x 2 contingency table, testing the significance of the observed intersection under random selection of two sets of 15 compounds from the full library of 527 drugs.

Associations between CES values and pharmacological metrics were assessed using Spearman rank correlation (ρ). In selected analyses evaluating agreement between matched datasets or compound profiles across models, Pearson correlation (r) was used to quantify linear concordance. Correlations were computed using pairwise complete observations when missing values were present. Boxplots display the median and interquartile range (IQR), and hierarchical clustering was performed using the Chebyshev distance metric with complete linkage. All reported *P* values are two-sided unless otherwise stated. Plate-level assay quality was assessed using standard screening metrics, including the Z′, robust Z′, signal-to-background ratio, and strictly standardized mean difference (SSMD) based on Breeze-QC procedures^56^. A complete summary of plate-level quality control statistics across all experimental conditions is provided in **Supplementary Data 2**.

### Computational pipeline for co-culture dose-response modeling

The pipeline was implemented through the following stages (see **Supplementary Figure 1**).

#### Stage 1. Quality control, data processing, and integrated co-culture profiling

Plate quality was assessed with standard metrics (Z-prime, robust Z-prime, signal-to-background ratio, and SSMD). Cell viability measurements were normalized to plate-specific controls to account for baseline effector-mediated target killing. Dose-response curves were independently fitted for each assay condition and interpolated onto a dense, uniform dose grid. Co-culture interaction profiles were then constructed according to the available experimental configuration and the selected scoring strategy (detailed in **Construction of integrated co-culture interaction profiles**). All constructed profiles were bounded between [−100, 100] to ensure numerical stability in downstream analyses.

#### Stage 2. Gaussian mixture modeling

Integrated co-culture profiles were modeled using a four-component Gaussian basis-function model to capture complex, non-linear response landscapes. Dose values were shifted to a non-negative domain to ensure optimization stability. Co-culture profiles were pre-processed so that (i) those with less than 10% absolute activity were assigned CES = 0, (ii) predominantly negative profiles were temporarily inverted to allow consistent parameter estimation, and (iii) mixed-sign profiles (minimum < -10%) were vertically shifted by their absolute minimum to maintain non-negativity. Model parameters were estimated by minimizing the sum of squared errors using bounded multi-start nonlinear optimization, with differential evolution as a global-search fallback.

#### Stage 3. Feature extraction and CES computation

The fitted Gaussian mixture model was used to reconstruct a continuous response curve from which pharmacological parameters were extracted: normalized area under the curve (nAUC), maximal efficacy of the response (Peak), and the corresponding effective concentration. Following back-correction for pre-processing shifts, these features were integrated into the Co-culture Efficacy Score, yielding a unified metric of compound activity and efficacy.

#### Stage 4. Automated output generation

The pipeline exports standardized datasets containing CES values, derived pharmacological features, and effector toxicity classifications. Interactive graphical outputs overlay condition-specific dose-response fits with the reconstructed co-culture interaction profile. Model diagnostics, including raw measurements, curve-fitting parameters (e.g., IC_50_, slope), and residual error statistics, were generated to support transparent data interpretation.

### Construction of integrated co-culture interaction profiles

To quantify context-dependent compound activity in co-culture assays, CES constructs a signed interaction profile from fitted dose-response curves evaluated on a common log_10_ concentration grid. Let R_CC_(c), R_T_(c), R_E_(c) denote the percent inhibition at concentration *c* in the co-culture, target-cell monoculture and effector-cell monoculture conditions, respectively. All responses are expressed on the same percent-inhibition scale after condition-specific normalization.

In the three-condition therapeutic mode, CES compares the observed co-culture response with the expected combined effect of direct target-cell inhibition and effector-cell toxicity under Bliss independence:

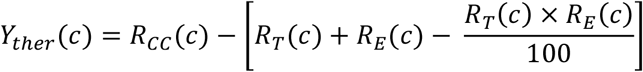

This mode prioritizes co-culture activity that exceeds the expected direct effects while penalizing compounds with direct effector-cell toxicity.

In the three-condition mechanistic mode, CES first computes the differential co-culture response relative to the target-cell monoculture and then scales this signal by the surviving effector-cell fraction:

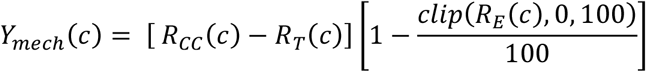

where *clip* (*R*_*E*_ (*c*), 0, 100) constrains the effector-cell inhibition value to the interval 0-100%. This mode attenuates interaction signals that are driven by effector-cell loss and avoids inflation from negative effector-cell inhibition values.

When only co-culture and matched reference conditions are available, CES is computed in two-condition mode:

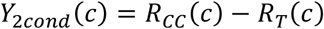

This mode quantifies differential co-culture activity relative to the matched reference condition but does not separately resolve effector-cell toxicity. The resulting interaction profile Y(c) is bounded to the interval [-100, 100] before Gaussian fitting. Positive values indicate increased co-culture activity relative to the reference condition, whereas negative values indicate reduced co-culture activity. In immune co-culture assays, positive values correspond to enhanced target-cell killing and negative values correspond to inhibition of target-cell killing. In host-pathogen rescue assays, the response orientation was set so that positive values correspond to host-cell rescue from virus-induced cytopathic effect.

### Gaussian fitting and CES calculation

Dose values were modeled on the log_10_ concentration scale, *x*_*i*_ = log_10_(*c*_*i*_ ), where *c*_*i*_ is the tested concentration. To improve numerical stability during optimization, the dose axis was shifted to a non-negative domain:

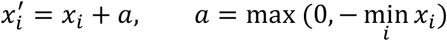

Here, 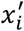 denotes the shifted log_10_ concentration used for model fitting, and *a* is the applied dose-axis shift. The tested log_10_ concentration range was defined as:

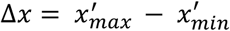

CES first applies activity and sign preprocessing to the interaction profile. Profiles with less than 10% absolute activity were considered inactive and assigned CES = 0. Predominantly negative profiles were multiplied by −1 before fitting and the sign was restored after feature extraction. Mixed-sign profiles with values below -10% were shifted upward by |min (*Y*)| before fitting and then back-corrected after model estimation. This preprocessing allows the Gaussian model to use non-negative component amplitudes while preserving the final direction and magnitude of the interaction profile.

The preprocessed interaction profile was summarized using a four-component Gaussian basis-function model:

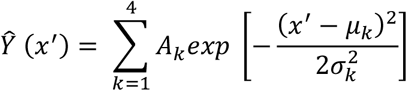

where *A*_*k*_, *µ*_*k*_ and *σ*_*k*_ denote the amplitude, center and standard deviation of Gaussian component *k*, respectively. The Gaussian model is used as a flexible smoothing function for summarizing the interaction profile; individual Gaussian components are not interpreted biologically.

Model parameters are estimated by minimizing the residual sum of squares:

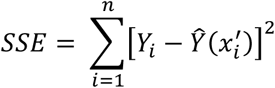

Optimization uses bounded multi-start nonlinear optimization with the nlminb function in R. Up to three starting configurations are evaluated, and the best solution is selected by minimum residual error. Evaluation terminates early if a configuration reaches a residual sum of squares that is negligible relative to the total variation of the interaction profile. To prevent degenerate narrow-peak solutions, each Gaussian component is constrained by a minimum standard deviation:

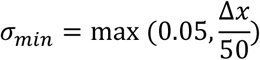

In case all local optimization attempts fail, differential evolution is used as a global-search fallback.

The analytic area under each Gaussian component over the tested concentration interval is computed as:

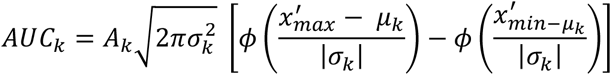

where *ϕ* denotes the standard normal cumulative distribution function. The normalized cumulative activity is calculated as:

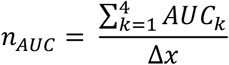

Here, the analytic areas of the four Gaussian components (*k* = 4) are summed and divided by the width of the tested concentration interval Δ*x* on the shifted log_10_ scale, yielding a cumulative activity measure normalized to the width of the tested concentration interval.

The peak effect is defined as the maximum value of the fitted profile over the tested concentration interval after the corresponding back-correction:

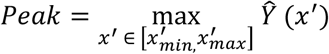

The reported peak effect is constrained to the interval [−100, 100]. The model-estimated effective concentration is defined as the original-scale concentration at which the fitted profile reached its maximum value:

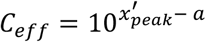

where 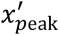 is the shifted log_10_ concentration at which the fitted profile has maximal value.

Finally, CES is calculated, with the scoring peak floored at 10 to keep the logarithm well-defined:

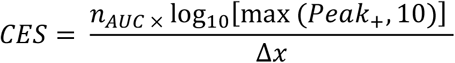

where Peak_+_ denotes the positive working-scale peak used for scoring prior to sign restoration of predominantly negative profiles. For negative profiles, the sign of CES is restored after calculation. This formulation combines normalized cumulative activity and maximal fitted effect while preserving the direction of the interaction profile. The model-estimated effective concentration is constrained to the tested concentration range.

## Software implementation

To facilitate broad accessibility and rapid deployment, the CES computational framework was deployed as an open-source, interactive web application (https://ces.aittokallio.group). This online resource automates the data processing pipeline, accepting both raw and pre-normalized multiplexed screening data as input. The backend executes the custom parametric Gaussian modeling, calculates condition-specific CES metrics, and applies the effector-toxicity penalties. The frontend interface provides interactive visualization modules for plate-level quality control (when plate annotation data is provided), dose-response curves per condition, toxicity analysis, and comprehensive compound prioritization by the CES features, allowing end-users to seamlessly perform high-dimensional screening analyses without requiring local bioinformatics infrastructure or programming experience.

## Code availability

The open-source R code for the CES algorithm is publicly available on GitHub (https://github.com/dias-dio/CES), with user instructions, enabling researchers to implement the methodology locally for custom co-culture drug screening pipelines. To maximize accessibility, the fully automated CES framework is also freely provided as an interactive web-tool (https://ces.aittokallio.group). Comprehensive user documentation, including required dataset structures, formatting guidelines, and step-by-step analysis tutorials, is provided both in the GitHub repository and on the dedicated documentation page of the online platform.

## Data availability

The raw and processed drug screening datasets generated and analyzed in this study are available in the Supplementary Data files. Specifically, drug-response measurements are provided in **Supplementary Data 1**, and the plate-level quality control parameters, Co-culture Efficacy Score metrics, and extended validation datasets are detailed in **Supplementary Data 2**-**9**. To facilitate interactive exploration, the sample datasets and application examples from the paper can be analyzed and visualized through the CES web-app at https://ces.aittokallio.group.

## Acknowledgments

We thank the Finnish Red Cross Blood Service for providing NK donor blood samples, and the healthy donors for their generous contributions. Drug sensitivity and resistance testing was performed at the FIMM High Throughput Biomedicine Unit, University of Helsinki, which provided the compound libraries, automated dispensing, and screening infrastructure used in this study; the unit is supported by HiLIFE and Biocenter Finland. We acknowledge the computing resources provided by CSC - IT Service for Science Ltd, and thank J.B. and FIMM colleagues for beta-testing the codes and web-application. This work was supported by the Jane and Aatos Erkko Foundation, the Research Council of Finland (grants 344698, 345803, 367855, and 373493 to T.A.), the Sigrid Juselius Foundation, the Signe and Ane Gyllenberg Foundation, the Cancer Foundation Finland, Blood Cancer United (formerly The Leukemia & Lymphoma Society), State funding for University-level Health Research in Finland, iCAN – Digital Precision Cancer Medicine Flagship, and HiLIFE Fellow funds. D.D. was supported by iCANDOC funding.

## Ethics declaration

Access to primary NK cells was approved by the Finnish Red Cross Biobank Ethics Committee. All donor-derived material was handled as sensitive and confidential data in accordance with applicable ethical and privacy regulations.

## Author Contributions

**D.D**. conceived the study, designed and developed the methodology and the interactive web application, performed the modeling and analysis of co-culture dose-response profiles, analyzed the experimental data, interpreted the results, prepared the figures, and drafted the manuscript. **A.I**. designed and developed the methodology and scoring equations, provided analytical interpretation of the results, and performed testing of the co-culture dose-response modeling framework. **J.B**. designed and conducted the experiments, pre-selected the compounds for the validation panel, performed the effector-to-target ratio optimization studies, and contributed to the development of the methodology. **O.D**. provided the CAR T-cell co-culture dataset and guidance on the biological interpretability of the results. **P.N**. conducted drug screening experiments. **J.K**. and **H.L**. conducted drug screening experiments, performed NK-cell expansions, and prepared luciferase-expressing cell lines. **J.B**. and **M.C**. extensively tested the interactive web application and provided feedback on the user experience and data visualization modules. **T.A**. and **S.M**. supervised the project, provided analytical interpretation of the results, and contributed to the critical revision and writing of the manuscript. All authors reviewed and approved the final version of the manuscript.

