## Supplementary Information for "Selective scoring of drug effects in multicellular co-culture systems"

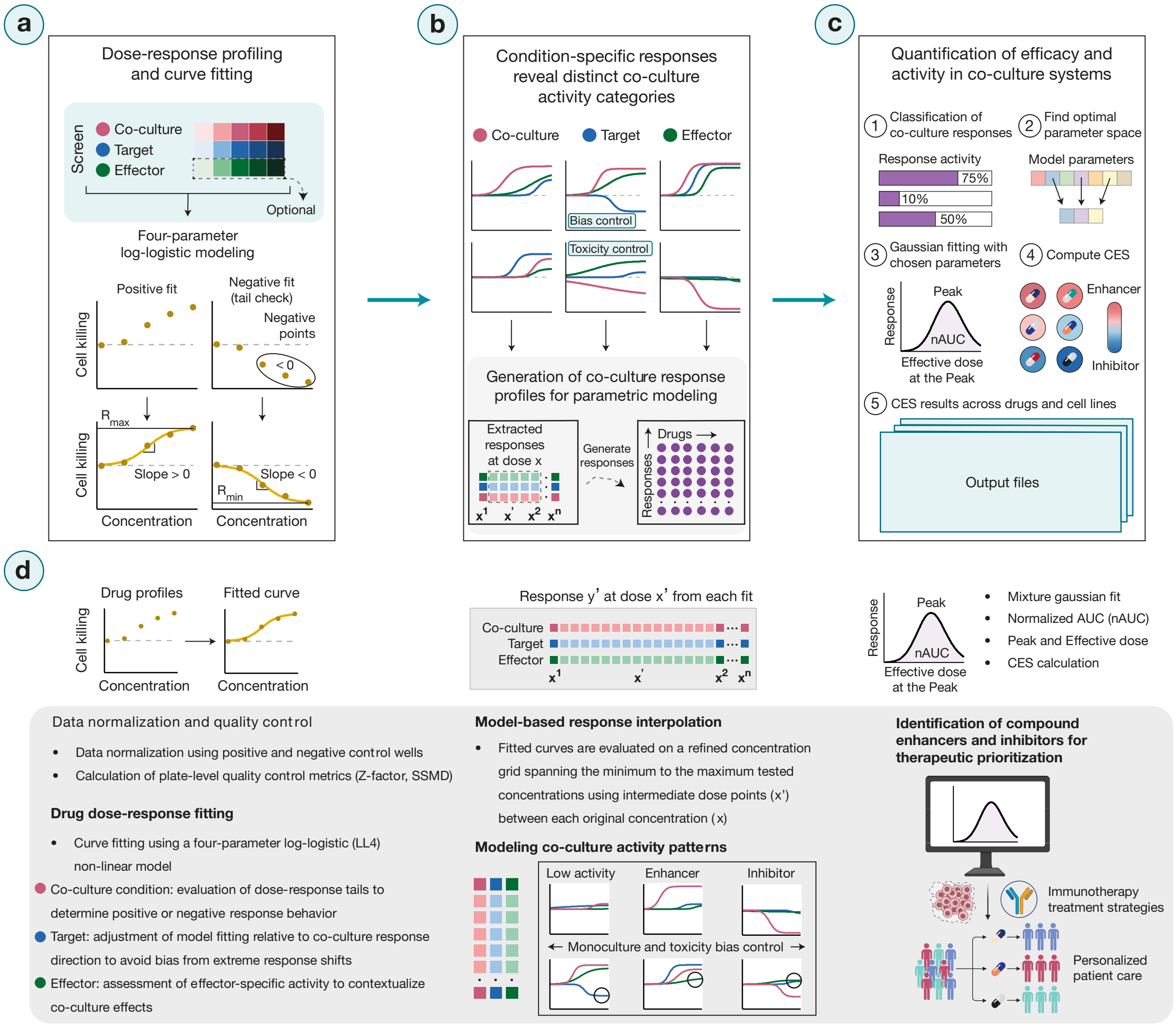


**Supplementary Figure 1. Technical workflow of co-culture dose-response modelling and CES computation.** (**a**) Dose-response measurements from high-throughput drug sensitivity screens are generated for three experimental conditions: target cell monoculture, target-effector co-culture, and effector cell monoculture. These data are fitted using four-parameter log-logistic modelling to generate individual dose-response curves. The curve-fitting process evaluates key response parameters, including slope, minimum and maximum responses ($R_{min}$ and $R_{max}$), and goodness of fit, to ensure valid response trajectories for downstream analysis. (**b**) Dose-response curves from the three experimental conditions are merged to derive co-culture response profiles. From these modelled curves, dose levels and corresponding response estimates are extracted to generate a unified representation of drug effects in the co-culture system. (**c**) Quantification of co-culture drug activity is performed using efficacy and cumulative activity parameters derived from the co-culture response profiles. The analysis includes (i) classification of co-culture response patterns, (ii) identification of an optimal parameter space for modelling, and (iii) Gaussian mixture modelling of the response profiles using the selected parameters. From the fitted curves, maximal drug effect (Peak) and cumulative activity (nAUC) are computed and integrated to derive the Co-culture Efficacy Score (CES). Standardized output files that summarize CES values across drugs and cell lines are generated for downstream analyses. (**d**) A summary of the analytical workflow corresponding to panels a–c, detailing dose-response modelling, data integration across experimental conditions, classification of co-culture activity patterns, and identification of compounds with enhanced or reduced activity in co-culture.

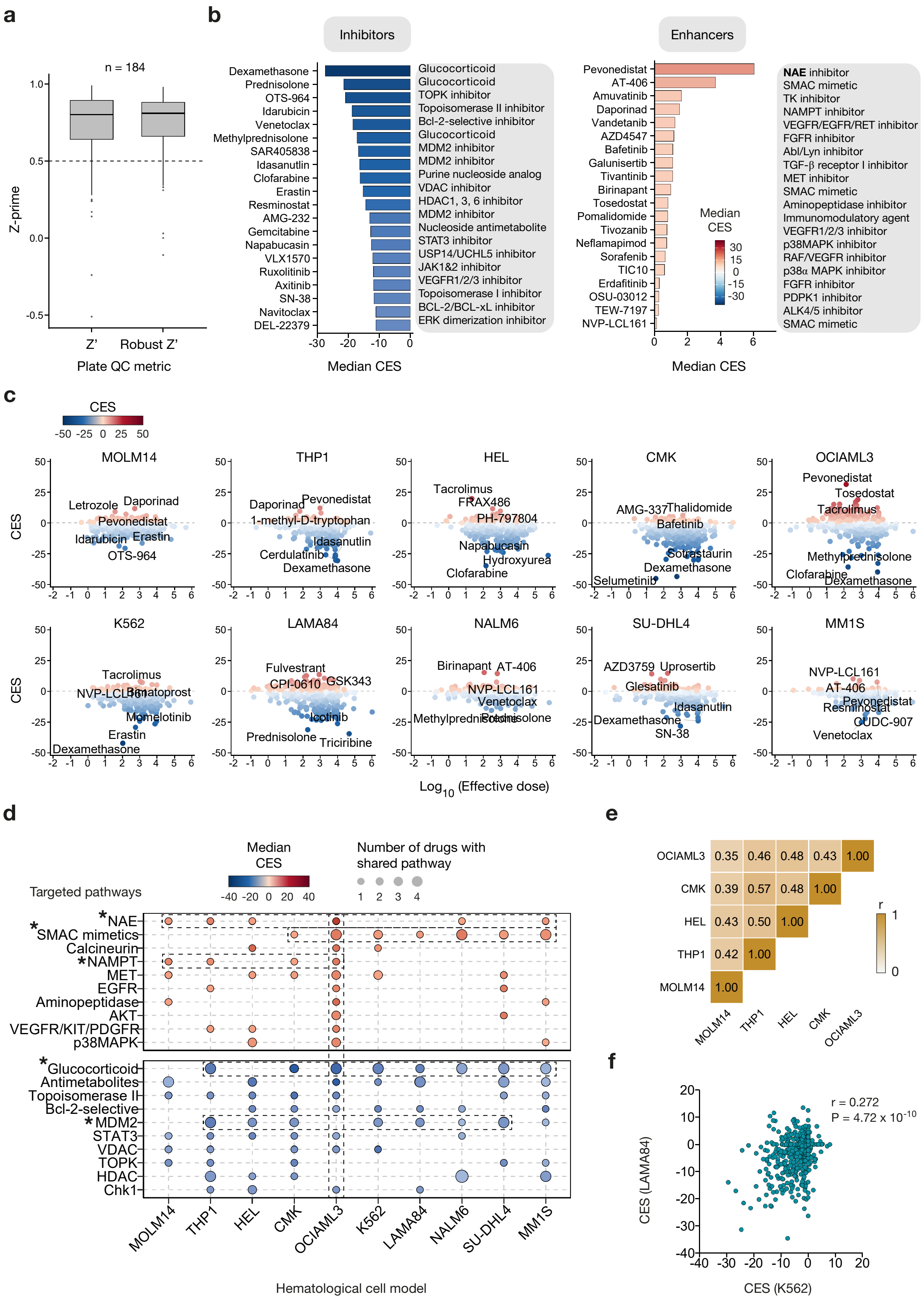
a
n = 184
Z-prime
Z'
Robust Z'
Plate QC metric
b
Inhibitors
Dexamethasone
Prednisolone
OTS-964
Idarubicin
Venetoclax
Methylprednisolone
SAR405838
Idasanutlin
Clofarabine
Erastin
Resminostat
AMG-232
Gemcitabine
Napabucasin
VLX1570
Ruxolitinib
Axitinib
SN-38
Navitoclax
DEL-22379
Glucocorticoid
Glucocorticoid
TOPK inhibitor
Topoisomerase II inhibitor
Bcl-2-selective inhibitor
Glucocorticoid
MDM2 inhibitor
MDM2 inhibitor
Purine nucleoside analog
VDAC inhibitor
HDAC1, 3, 6 inhibitor
MDM2 inhibitor
Nucleoside antimetabolite
STAT3 inhibitor
USP14/UCHL5 inhibitor
JAK1&2 inhibitor
VEGFR1/2/3 inhibitor
Topoisomerase I inhibitor
BCL-2/BCL-xL inhibitor
ERK dimerization inhibitor
Median CES
Enhancers
Pevonedistat
AT-406
Amuvatinib
Daporinad
Vandetanib
AZD4547
Bafetinib
Galunisertib
Tivantinib
Birinapant
Tosedostat
Pomalidomide
Tivozanib
Neflamapimod
Sorafenib
TIC10
Erdafitinib
OSU-03012
TEW-7197
NVP-LCL161
NAE inhibitor
SMAC mimetic
TK inhibitor
NAMPT inhibitor
VEGFR/EGFR/RET inhibitor
FGFR inhibitor
Abl/Lyn inhibitor
TGF-β receptor I inhibitor
MET inhibitor
SMAC mimetic
Aminopeptidase inhibitor
Immunomodulatory agent
VEGFR1/2/3 inhibitor
p38MAPK inhibitor
RAF/VEGFR inhibitor
p38α MAPK inhibitor
FGFR inhibitor
PDPK1 inhibitor
ALK4/5 inhibitor
SMAC mimetic
Median CES
Median CES
c
CES
MOLM14
THP1
HEL
CMK
OCIAML3
K562
LAMA84
NALM6
SU-DHL4
MM1S
Log10 (Effective dose)
d
Targeted pathways
Median CES
Number of drugs with shared pathway
*NAE
*SMAC mimetics
Calcineurin
*NAMPT
MET
EGFR
Aminopeptidase
AKT
VEGFR/KIT/PDGFR
p38MAPK
*Glucocorticoid
Antimetabolites
Topoisomerase II
Bcl-2-selective
*MDM2
STAT3
VDAC
TOPK
HDAC
Chk1
Hematological cell model
e
r
f
r = 0.272
P = 4.72 x 10-10
CES (LAMA84)
CES (K562)

**Supplementary Figure 2**. **Quality control and cross-model relationships of CES values across hematological cancer cell lines**. (**a**) Distribution of plate-level quality control metrics across all screening plates. Boxplots show Z′ and robust Z′ statistics computed from positive and negative control wells for each assay plate (*n* = 184). Both metrics indicate consistent assay performance across co-culture, target monoculture, and effector monoculture conditions. (**b**) Barplots show the top 20 CES-enhancing and top 20 CES-inhibitory compounds ranked by median CES across the ten hematological cancer cell lines. Bars represent the median CES value per drug across cell lines. Positive values (red gradient) indicate enhancement of effector-mediated cytotoxicity, whereas negative values (blue gradient) indicate inhibition. Drugs are ordered by magnitude of median CES within each group. (**c**) Scatterplots of CES against -$\log_{10}$ transformed effective concentrations for each hematological cancer cell line. Each point represents one drug (a total of 527 per panel). Point colour indicates CES using a global symmetric red-blue scale shared across panels. Horizontal dashed lines indicate CES = 0, separating enhancers from inhibitors. The three strongest enhancers and inhibitors per cell line are annotated. Axes and colour scales are held constant across panels to enable direct comparison between models. (**d**) Pathway-level summary of enhancer (top) and inhibitor (bottom) compounds across hematological cancer models. For each cell line, compounds were grouped by pharmacological class and the median therapeutic CES computed per class. The 10 pathway classes with the strongest total signal are shown. Dot color indicates CES within each cell line-pathway pair and dot size reflects the number of contributing compounds. (**e**) Upper-triangular pairwise Spearman correlation matrix of CES values across AML cell line models. Correlations were computed across all screened compounds using pairwise complete observations (see **Methods**). (**f**) Correlation of CES values between the two CML cell lines. Each point represents one compound profiled in both K562 and LAMA84 models. Spearman correlation coefficient (ρ) quantifies concordance of CES responses between the two CML models.

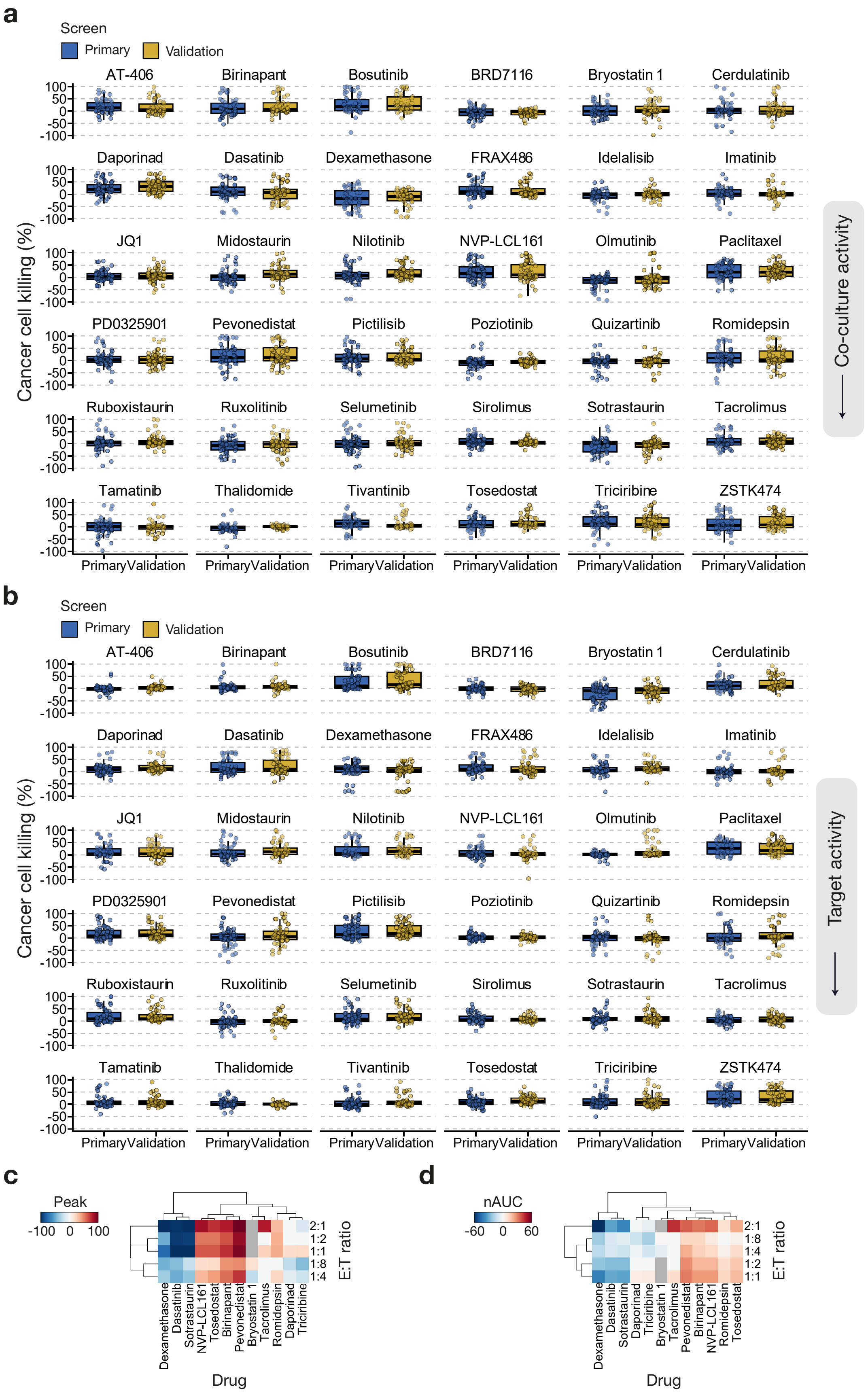
a
Screen
Primary
Validation
AT-406
Birinapant
Bosutinib
BRD7116
Bryostatin 1
Cerdulatinib
Daporinad
Dasatinib
Dexamethasone
FRAX486
Idelalisib
Imatinib
JQ1
Midostaurin
Nilotinib
NVP-LCL161
Olmutinib
Paclitaxel
PD0325901
Pevonedistat
Pictilisib
Poziotinib
Quizartinib
Romidepsin
Ruboxistaurin
Ruxolitinib
Selumetinib
Sirolimus
Sotrastaurin
Tacrolimus
Tamatinib
Thalidomide
Tivantinib
Tosedostat
Triciribine
ZSTK474
100
50
0
-50
-100
Cancer cell killing (%)
Primary Validation
Co-culture activity
b
Screen
Primary
Validation
Cancer cell killing (%)
Target activity
c
Peak
-100 0 100
2:1
1:2
1:1
1:8
1:4
E:T ratio
Dexamethasone
Dasatinib
Sotrastaurin
NVP-LCL161
Tosedostat
Birinapant
Pevonedistat
Bryostatin 1
Tacrolimus
Romidepsin
Daporinad
Triciribine
Drug
d
nAUC
-60 0 60
2:1
1:8
1:4
1:2
1:1
E:T ratio
Dexamethasone
Dasatinib
Sotrastaurin
Daporinad
Triciribine
Bryostatin 1
Tacrolimus
Pevonedistat
Birinapant
NVP-LCL161
Romidepsin
Tosedostat
Drug

**Supplementary Figure 3**. **Comparison of primary and validation screens and drug response relationships across E:T ratios**. (**a**) Percent inhibition of matched compounds in the primary and validation co-culture screens. Analysis was restricted to overlapping drug concentrations to ensure direct comparability. Individual points represent measurements from a given cell line at a given concentration. Boxplots show the median and interquartile range (IQR). (**b**) Percent inhibition of matched compounds in the primary and validation target screens, shown as in panel a. Statistical significance was evaluated for each compound using a two-sided Wilcoxon signed-rank test across the ten cell lines, followed by Benjamini–Hochberg (BH) correction for multiple testing; no significant differences were observed between the primary and validation screens for any compound (all adjusted $P > 0.05$). (**c**) Hierarchical clustering heatmap of maximal effect (Peak) values across E:T ratios for the 12 selected compounds in the OCIAML3 model. Rows represent E:T conditions and columns represent compounds. Distances were computed using the Chebyshev metric with complete linkage. Colour intensity reflects deviation from zero-centered Peak values. (**d**) Hierarchical clustering heatmap of cumulative activity (nAUC) values across E:T ratios for the same compound set, with clustering and colour scaling performed analogously to panel c.

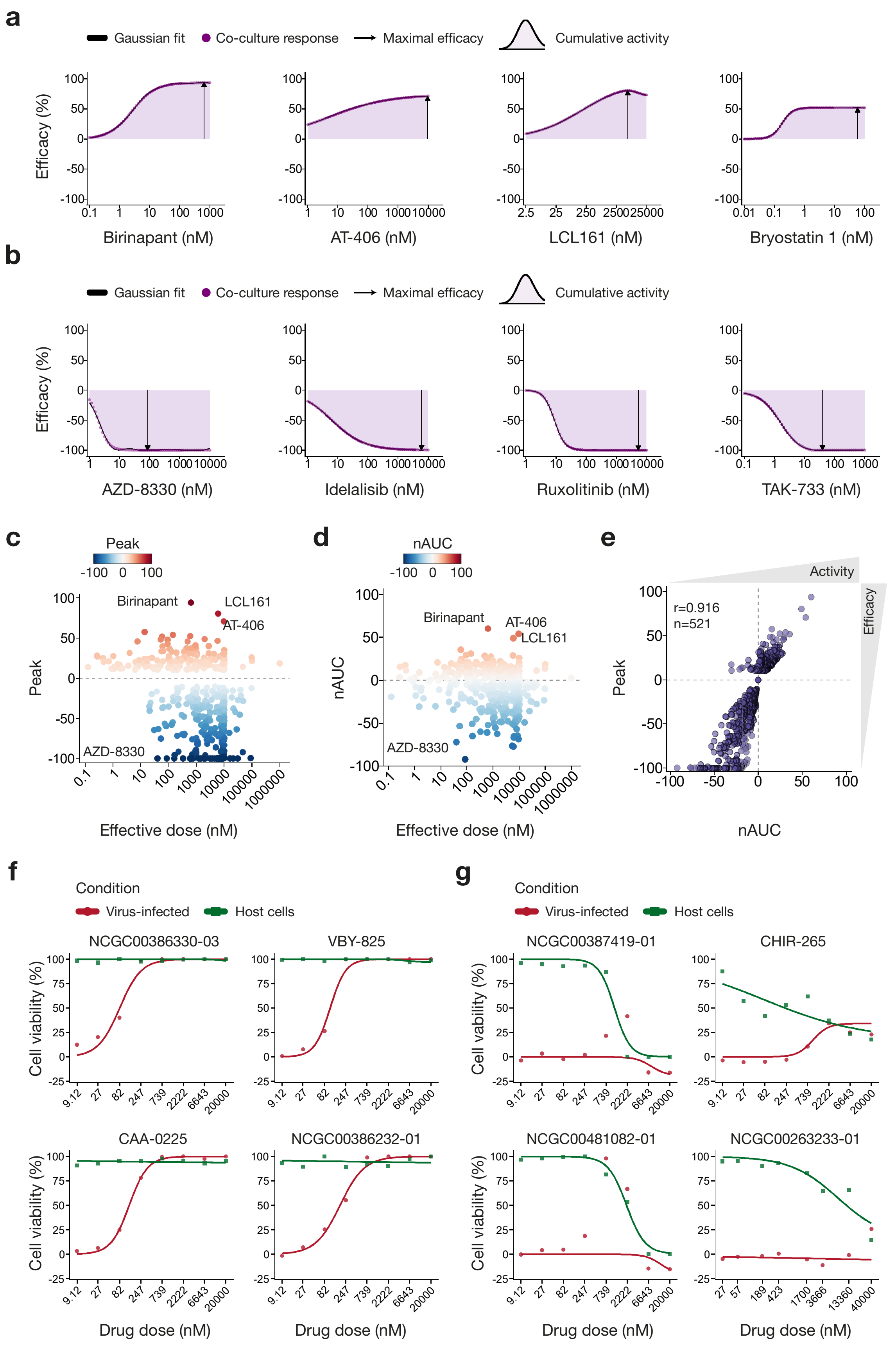
a
Gaussian fit
Co-culture response
Maximal efficacy
Cumulative activity
Efficacy (%)
Birinapant (nM)
AT-406 (nM)
LCL161 (nM)
Bryostatin 1 (nM)
b
AZD-8330 (nM)
Idelalisib (nM)
Ruxolitinib (nM)
TAK-733 (nM)
c
Peak
Birinapant
LCL161
AT-406
AZD-8330
Effective dose (nM)
d
nAUC
e
Activity
Efficacy
r=0.916
n=521
f
Condition
Virus-infected
Host cells
NCGC00386330-03
VBY-825
CAA-0225
NCGC00386232-01
Cell viability (%)
Drug dose (nM)
g
NCGC00387419-01
CHIR-265
NCGC00481082-01
NCGC00263233-01

**Supplementary Figure 4. Dose-response curves and CES metrics for the CAR T-cell co-culture and antiviral screening datasets.** (**a**) Modelled co-culture dose-response curves for the four compounds exhibiting the highest positive CES values (enhancers) in the CAR T-cell co-culture assay. (**b**) Modelled co-culture dose-response curves for the four compounds demonstrating the most negative CES values (inhibitors) in the CAR T-cell co-culture assay. (**c**) Maximal effect (Peak) plotted against effective dose across compounds in the CAR T-cell co-culture dataset. Effective dose is displayed on a $-\log_{10}$ scale (nM), and points are coloured by Peak value. (**d**) Cumulative activity (nAUC) plotted against effective dose across 527 compounds in the CAR T-cell co-culture dataset; axes as in panel (c), with points coloured by nAUC value. (**e**) Relationship between efficacy (Peak) and cumulative activity (nAUC) in the CD19-targeting CAR T-cell co-culture dataset. (**f**) Fitted dose-response curves for the four compounds with the highest positive CES values in the antiviral screening dataset. The x-axis represents drug concentration (nM) and the y-axis represents viability response (host-cell activity post-infection). (**g**) Fitted dose-response curves for the four compounds with the most negative CES values in the antiviral screening dataset; axes as in panel f.

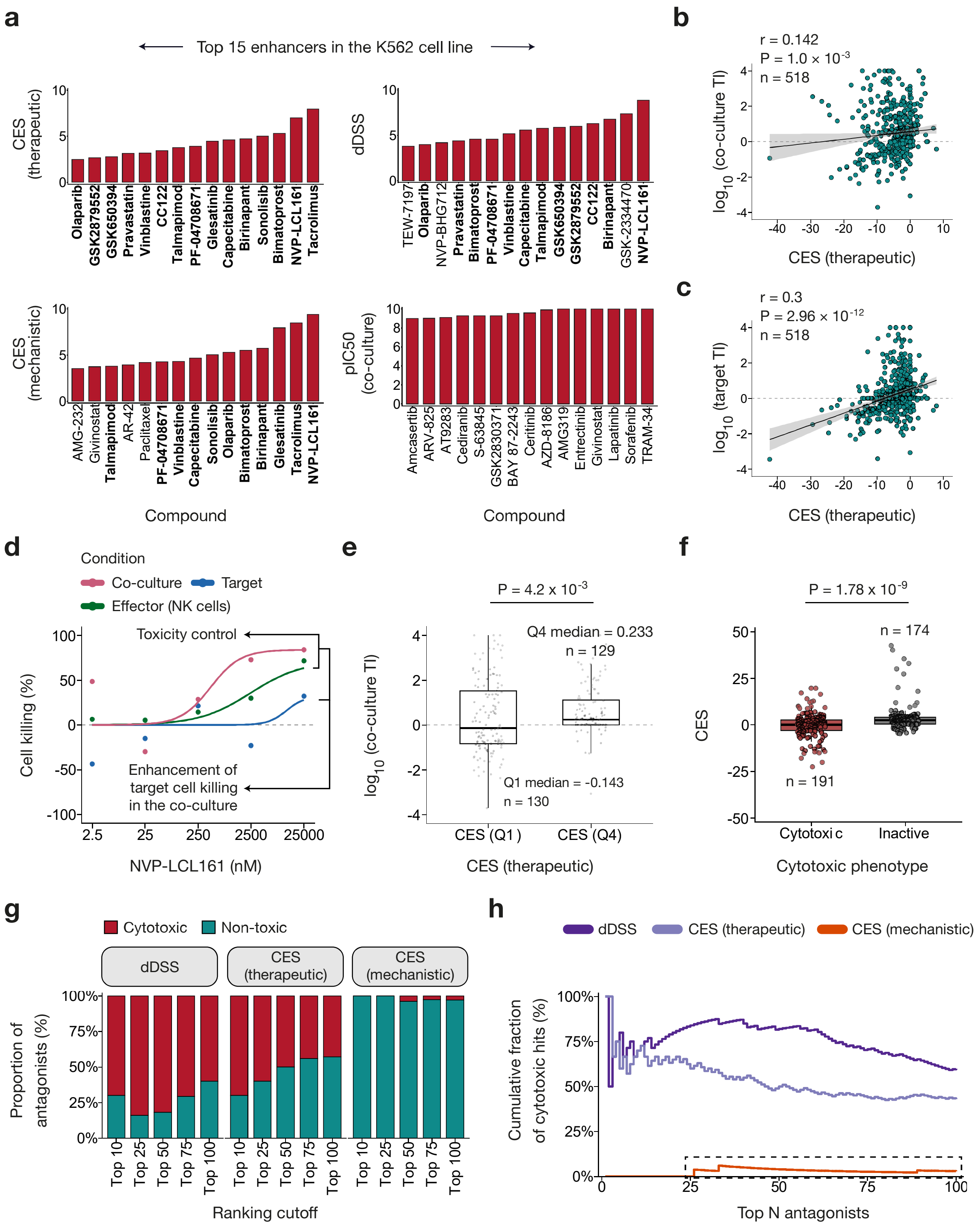


**Supplementary Figure 5**. **Extended analysis of CES resolution across immune and antiviral contexts**. (**a**) Top 15 enhancer compounds in the K562 NK-cell co-culture screen (a total of 527 compounds) ranked by therapeutic CES, mechanistic CES, dDSS, and co-culture $pIC_{50}$ ($-log_{10}$ of the fitted $IC_{50}$ in molar units). As in Fig. 5b, $IC_{50}$ is included as the conventional single-curve baseline, whereas therapeutic CES, mechanistic CES, and dDSS integrate information across full dose-response profiles and reference conditions. Compounds shared between therapeutic CES and the alternative scoring metrics are emphasized with bold text. (**b**) Association between therapeutic CES and the co-culture therapeutic index (TI), defined as $\log_{10}$(NK-cell $IC_{50}$ / co-culture $IC_{50}$), in the K562 screen (518 compounds with finite TI). The Spearman correlation coefficient (ρ) and associated P-value are reported. The dashed line indicates $\log_{10}TI = 0$. (**c**) Association between therapeutic CES and the target-cell monoculture therapeutic index (TI), defined as $\log_{10}$(NK-cell $IC_{50}$ / target-cell $IC_{50}$), in the K562 screen (518 compounds with finite TI). The Spearman correlation coefficient (ρ) and associated P-value are reported. The dashed line indicates $\log_{10}TI = 0$. (**d**) Dose-response curves for NVP-LCL161, the consistently top-ranked enhancer shared across therapeutic CES, mechanistic CES, and dDSS, in the K562 co-culture screen, with the three conditions shown. Arrows indicate how CES controls for direct effector-cell toxicity when quantifying the enhancement of target-cell killing in co-culture. (**e**) Co-culture therapeutic index (TI) values, defined as $\log_{10}$(NK-cell $IC_{50}$ / co-culture $IC_{50}$), for compounds in the K562 screen in the bottom (Q1) and top (Q4) quartiles of therapeutic CES (n = 130 and 129 compounds, respectively). Boxes show median and interquartile range (IQR); the dashed line indicates $\log_{10}TI = 0$. Differences were tested with a two-sided Wilcoxon rank-sum test. (**f**) Distribution of CES values stratified by antiviral control-assay phenotype (cytotoxic vs inactive). Differences were tested with a two-sided Wilcoxon rank-sum test. (**g**) Composition of the top N antagonists by toxicity for each scoring metric in the K562 screen, shown at multiple ranking cutoffs of N = 10, 25, 50, 75, and 100. Stacked bars display the proportion of broadly cytotoxic compounds (red) versus non-toxic antagonists (teal) prioritized by dDSS, therapeutic CES, and mechanistic CES. Compounds were classified as cytotoxic once the drug sensitivity score against the NK-cell control condition exceeded the threshold (DSS ≥ 10). (**h**) Cumulative fraction of broadly cytotoxic compounds (DSS ≥ 10 of the effector cells) among the top N predicted antagonists for each scoring metric in the K562 screen, evaluated continuously from the top 1-100 antagonists. Mechanistic CES identified on average 2.6% cytotoxic compounds across the top 100 antagonists, compared with 53.4% for therapeutic CES and 75.7% for dDSS.

**Supplementary Table 1. Summary statistics of therapeutic CES across 10 hematological cancer cell lines co-cultured with NK cells and screened against 527 compounds.**

| Disease | Cell line | Drugs | Enhancers | Inhibitors | Mean | Median | Sd | Min | Max |
|---|---|---|---|---|---|---|---|---|---|
| AML | MOLM14 | 515 | 81 | 346 | -2.59 | -1.95 | 4.00 | -20.5 | 11.6 |
| AML | THP1 | 512 | 59 | 351 | -3.91 | -2.51 | 5.58 | -30.6 | 12.1 |
| AML | HEL | 509 | 64 | 351 | -4.15 | -2.75 | 5.87 | -34.8 | 19.8 |
| AML | CMK | 516 | 54 | **408** | **-6.12** | **-4.87** | 6.62 | **-45.1** | 6.25 |
| AML | OCIAML3 | 521 | 118 | 322 | -2.00 | -1.44 | **6.79** | -39.8 | **31.4** |
| CML | K562 | 518 | 46 | 398 | -4.44 | -3.06 | 5.46 | -42.3 | 7.97 |
| CML | LAMA84 | 512 | **131** | 310 | -3.35 | -1.82 | 6.73 | -34.6 | 13.5 |
| B-ALL | NALM6 | 517 | 84 | 321 | -1.81 | -1.28 | 3.30 | -15.6 | 15.1 |
| BCL | SU-DHL4 | 516 | 58 | 331 | -2.70 | -1.36 | 4.65 | -28.5 | 14.5 |
| MM | MM1S | 509 | 38 | 269 | -1.68 | -0.51 | 3.37 | -25.3 | 9.99 |

For each cell model, the table reports the total number of evaluated compounds, the number of enhancer compounds (CES > 0), the number of inhibitor compounds (CES < 0), and descriptive statistics of the CES values. Differences in CES distributions across cell lines were assessed using a Kruskal-Wallis test ($P < 0.00001$). Boldfaced values indicate the most extreme values.

**Supplementary Table 2**. **Most frequent drug target classes among the top enhancer and inhibitor compounds across hematological cancer models.**

| Cell line | Disease type | Enhancer pathways | Inhibitor pathways |
|---|---|---|---|
| MOLM14 | AML | NAMPT inhibitor; Aromatase inhibitor; NAE inhibitor | TOPK inhibitor; Topoisomerase II inhibitor; VDAC inhibitor |
| THP1 | AML | NAE inhibitor; NAMPT inhibitor; IDO1/2 inhibitor | Glucocorticoid; MDM2 inhibitor; JAK, SYK inhibitor |
| HEL | AML | p38MAPK inhibitor; Calcineurin inhibitor; PAK1, 2, 3 inhibitor | Antimetabolite; Antineoplastic agent; CSC inhibitor, STAT3 mediated |
| CMK | AML | Immunosuppressant; MET inhibitor; Abl, Lyn inhibitor | Glucocorticoid; MEK1/2 inhibitor; PKC inhibitor |
| OCIAML3 | AML | SMAC mimetic; NAE inhibitor; Aminopeptidase inhibitor | Glucocorticoid; Antimetabolite; JAK, SYK inhibitor |
| K562 | CML | SMAC mimetic; Calcineurin inhibitor; Prostaglandin analog | Glucocorticoid; VDAC inhibitor, JAK 1 and 2 inhibitor |
| LAMA84 | CML | BET family inhibitor; EZH2 inhibitor; Estrogen receptor antagonist | Glucocorticoid; AKT inhibitor, EGFR inhibitor |
| NALM6 | B-ALL | SMAC mimetic; MCT1 inhibitor; Syk inhibitor | Glucocorticoid; Bcl-2-selective inhibitor; CDK7 inhibitor |
| SU-DHL4 | BCL | AKT inhibitor; EGFR inhibitor; MET/AXL/TIE/VEGFR, and RON inhibitor | Glucocorticoid; MDM2 inhibitor; Topoisomerase I inhibitor |
| MM1S | MM | SMAC mimetic; NAE inhibitor; Aminopeptidase inhibitor | HDAC inhibitor; Bcl-2-selective inhibitor; HDAC1/2/3/10 and PI3Kalpha inhibitor |

For each cell line model, the ten most enhancing and ten most inhibiting compounds were identified by therapeutic CES. Drug target annotations were harmonized into standardized mechanism-of-action labels, and the three most frequently represented classes in each category are reported. Cell lines are grouped by disease subtype. Full compound target annotations and counts are provided in **Supplementary Data 4**.

**Supplementary Data** (separate Excel files)

**Supplementary Data 1.** Dose-response percent inhibition measurements from the primary co-culture drug screen across the ten hematological cancer cell lines.

**Supplementary Data 2.** Plate-level quality control statistics for all screening plates across target monoculture, effector-cell monoculture, and co-culture conditions.

**Supplementary Data 3.** Therapeutic Co-culture Efficacy Score (CES) results and comprehensive dose-response metrics across the ten hematological cancer cell lines.

**Supplementary Data 4.** Pathway-level summary of representative compounds prioritized by therapeutic CES, reporting average efficacy and phenotypic classifications aggregated by hematological cancer subtype and cell line.

**Supplementary Data 5.** CES results from the independent validation experiment profiling a 36-compound panel at high dose resolution across the hematological cancer cell lines.

**Supplementary Data 6.** Therapeutic CES results across the five effector-to-target (E:T) ratios in the OCI-AML3 cell line, assessing metric robustness across varying effector loads.

**Supplementary Data 7.** CES results from the CAR T-cell co-culture screen against the CD19-positive NALM6 target cell line.

**Supplementary Data 8.** CES results, condition-specific dose-response parameters, and phenotypic classifications from the antiviral host-pathogen screening dataset.

**Supplementary Data 9.** Therapeutic and mechanistic CES profiles in the K562 hematological cancer cell line.
